# A Community Standard Multispecies Cell Atlas of the Basal Ganglia

**DOI:** 10.64898/2026.04.14.717814

**Authors:** The BRAIN Initiative Cell Atlas Network (BICAN), Joseph R. Ecker, Michael Hawrylycz, Ed Lein, Bing Ren, Carol Thompson, Hongkui Zeng, Owen White, Guo-Qiang Zhang

**Affiliations:** Howard Hughes Medical Institute, The Salk Institute for Biological Studies; Allen Institute for Brain Science; Allen Institute; New York Genome Center; Allen Institute, Brain Science; Univ. of Maryland; The University of Texas Health Science Center at Houston

**Author notes:** Correspondence: Joseph R. Ecker, Michael Hawrylycz, Ed Lein, Bing Ren, Carol Thompson, Hongkui Zeng, Owen White, Guo-Qiang Zhang. Author contributions at the end of the paper (The BRAIN Initiative Cell Atlas Network (BICAN)).

## Abstract

The NIH BRAIN Initiative Cell Atlas Network (BICAN) aims to generate a standardized, integrated cell atlas of the human, macaque, marmoset, and mouse brain that serves as a foundational community reference for the classification and study of brain cell types. Here we present the first major component of this effort: a cross-species, multimodal atlas of the basal ganglia, a group of subcortical nuclei central to motor control and implicated in a broad range of neurological disorders. Grounded in large-scale single-cell transcriptomic classification and integrated with epigenomic and spatial genomic modalities, this resource is enabled by coordinated cross-species sampling and harmonized analytical frameworks. It provides extensive phenotypic characterization of cell types, incorporates community-informed annotation, and establishes a highly curated, data-driven taxonomy with standardized nomenclature. The atlas is anchored to species-specific anatomical reference frameworks and linked across species through unified structural ontologies, enabling consistent cross-species comparisons. Multiple complementary datasets are mapped to this reference, including multiomic profiles and developmental trajectories aligned to adult cell states. Realization of this resource has required coordinated standards for tissue processing across human and model organisms, harmonization of donor metadata across brain banks, and the development of unified anatomical reference systems. To support these advances, BICAN has established an integrated ecosystem comprising standardized sequencing pipelines, neuroanatomically grounded data infrastructure, scalable visualization and mapping tools, and interoperable metadata standards. Analogous to the standardization achieved in genome science, this ecosystem provides a FAIR (findable, accessible, interoperable, and reusable) framework that enables researchers to map, compare, and interpret diverse datasets against a shared reference and associated knowledge base. The BICAN reference system is now being extended to the whole brain, with principles that are readily generalizable to other organ systems.

## 1. The BRAIN Initiative Cell Atlas Network

The mammalian brain is an extraordinarily complex organ, distinguished by immense cellular diversity, intricate local and long-range circuitry, and multiple modes of neural signaling^1^. The dynamic activity of neurons gives rise to complex adaptive behaviors and internal states, while also regulating essential physiological functions.^2,3^ These processes are mediated by highly specialized yet tightly integrated neural circuits composed of diverse cell types with distinct molecular, anatomical, and physiological properties.^4,5^ Understanding how brain function emerges from this diversity requires comprehensive knowledge of the cell types and circuits that together define the brain’s molecular and anatomical architecture.^5,6^ Accomplishing this at the scale of the human brain is a formidable challenge, and it therefore demands standardized, reproducible, and robust frameworks for describing cell types and their anatomical context.^6,7^ To address this need, the *National Institutes of Health (NIH) BRAIN Initiative Cell Atlas Network (BICAN)^8^*, a consortium of neuroscientists, computational biologists, and engineers, has been generating standardized and integrated cell atlases of the human, macaque, marmoset, and mouse brains. These atlases are intended to serve as community standards and versioned reference frameworks for mammalian brain organization.

The rapid adoption of single-cell molecular technologies has transformed our understanding of the diversity and complexity of brain cell types.^1,9,10^ Atlases have now been generated across multiple species, brain regions, and disease states, providing invaluable resources for studying cell types and their perturbations.^11–15^ Central to this enterprise is the development of reference frameworks for classifying the structure and variation of brain cell types^16^. When rigorously constructed, these atlases provide molecular reference standards analogous to representative genome references assembled from multiple individuals, thereby establishing a baseline against which variation can be measured^17^. Conceptually, cell type atlases resemble pangenomes in aggregating population-level variation into a shared reference^18^. However, whereas pangenomes encode largely discrete sequence variation, cell atlases must represent a high-dimensional and continuous cellular state space of the mammalian brain, in which discrete cell types coexist with graded, sex- and context-dependent variation.

A major challenge in studying the human brain is bridging contributions from diverse scientific fields and spatial scales from histological, cellular and molecular analyses to functional MRI (Magnetic Resonance Imaging) and macroscale connectomics.^19,20^ This challenge is compounded by the sheer scale of the human brain, which contains billions of neurons and glial cells organized into complex, distributed circuits, making comprehensive cellular characterization inherently difficult. Bridging these domains and integrating across scale is essential to constructing a transformative new cell atlas that links cellular and molecular features to the functional organization of the human brain^11^.

The complexity, diversity, and scale of brain cell types makes a unifying framework essential for classification and the investigation of function.^1,16^ Central to this framework is the classification of cell types and the ability to describe and map new data with respect to the reference^21^. Single-cell genomic analyses have revealed that brain cell types are highly conserved across mammals from mice to humans, providing a powerful framework for understanding molecular homology as well as species-specific differences.^12^ Cell type taxonomies can be aligned across species^12^ and, analogously to alignment of genome sequences, are more similar among evolutionarily closer primates^12^. This cross-species correspondence amplifies the value of non-human primate models for interpreting human brain organization and inferring cellular properties not directly accessible in human studies. Species-specific differences observed in jointly mapping these data are leading to the identification of uniquely human genetic features.^22–24^

The BICAN consortium is producing comprehensive cell atlases of human and non-human primate (NHP) brains that aspire to become new community standards and serve as foundational references for the neuroscience community. BICAN atlases leverage and integrate whole-brain taxonomies of the mouse brain produced in the BRAIN Initiative Cell Census Network (BICCN).^13,25–28^ Together, these atlases will serve as a key integrative reference connecting major brain initiatives and consortia, including the BRAIN Initiative Connectivity Across Scales (CONNECTS^29^), BRAIN Armamentarium^29,30^, the Human Cell Atlas (HCA^29–31^), and the HumanBioMolecular Atlas Program (HuBMAP^32^). The atlases will serve the neuroimaging community including the Human Connectome Project (HCP^32,33^), functional studies in brain circuitry, and many translational initiatives such as PsychENCODE^34^, Seattle Alzheimer’s Disease Brain Cell Atlas (SEA-AD ^35^), Michael J. Fox Foundation^34,36^, and Human Tumor Atlas Network^37^(**Fig. 1A**). BICAN atlases are creating fundamental knowledge on diverse cell types and their organizational principles in the brain across human, macaque, marmoset, and mouse (**Fig. 1B**), and across development and the lifespan^38^ of humans and mice (**Fig. 1C**). The outcome will produce a new reference classification for cell types and their epigenomic regulation across the whole human and NHP brain, spatial maps of molecularly defined cell types, and phenotypic characterization of fundamental brain cell types. The classification will also align homologous cell types from mice, marmosets, macaques and humans, allowing inference and comparison of cellular properties across species. Finally, data will be aligned in spatial common coordinate frameworks^38–40^, allowing creation of new atlases spanning structural, functional, cellular and molecular information.

**Figure 1.**
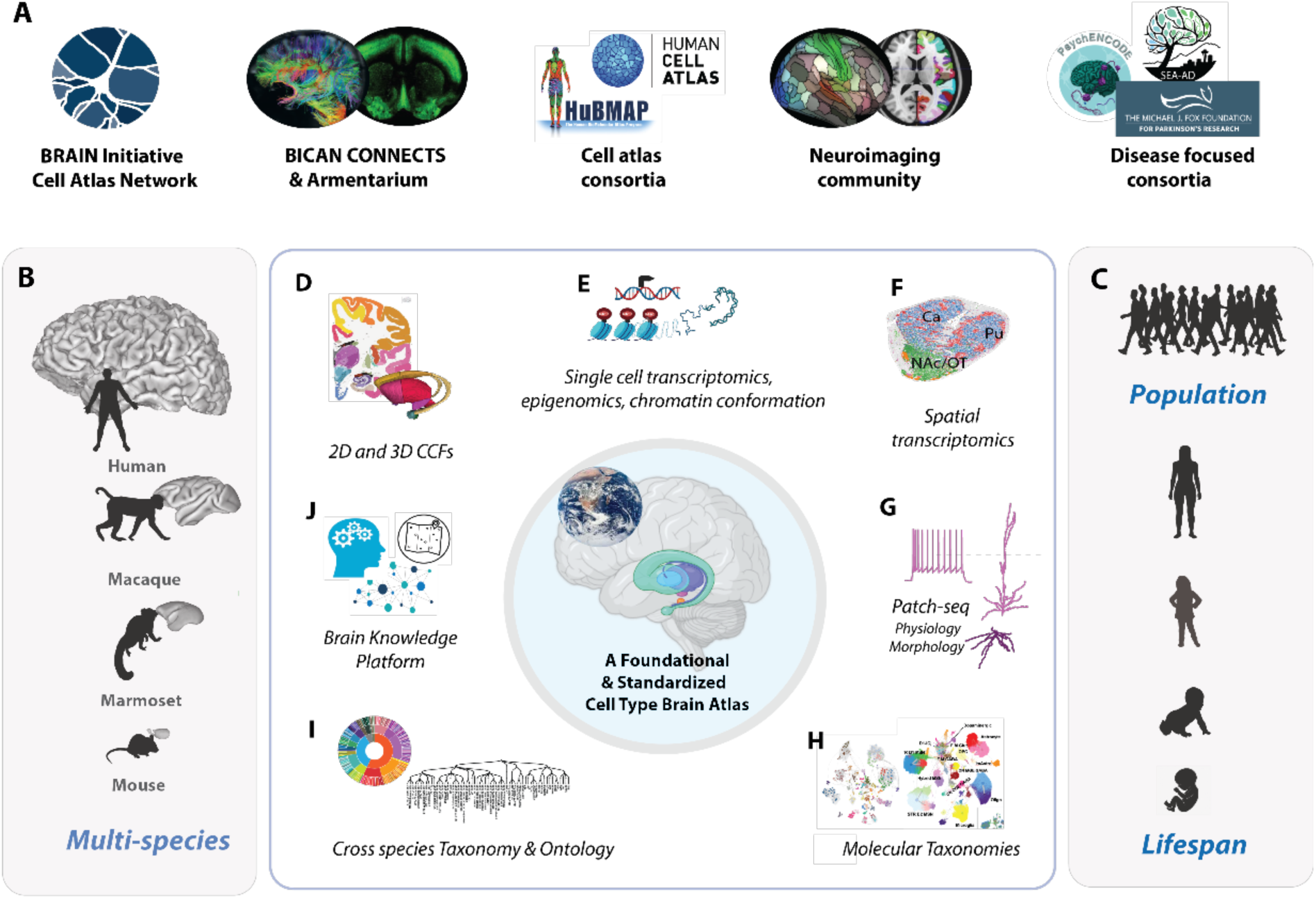
A Foundational Basal Ganglia Cell Type Atlas. A) BICAN is developing a foundational and standardized cell type atlas of the human and non-human primate brain, extending whole brain atlases of the mouse. The impact of this resource provides a cell type reference for connectivity studies, cell atlas consortia, the neuroimaging community, and disease-focused consortia. B) Multi-species brains shown to relative size. C) Atlasing across lifespan and population. A foundational atlas consists of several components: D) Common coordinate frameworks in 2D and 3D for data alignment and visualization. E-H) Multimodal data profiling with transcriptomic reference. I) Within species and consensus standardized molecular taxonomy and ontology. J) Brain Knowledge Platform, including access to publicly available data, data visualization, and analysis tools.

The basal ganglia (BG), comprising roughly 200 million neurons in the human brain, orchestrates the integration of cortical, thalamic, and brainstem signals to shape adaptive behaviors.^41^ Dysfunction within these circuits contributes to a wide spectrum of neurological and psychiatric disorders, including Parkinson’s disease, Huntington’s disease, dystonia, obsessive–compulsive disorder, and substance use disorders.^41–43^ In addition to their importance in translational applications, the BICAN consortium selected these structures first because of their strong evolutionary homology across species. The first component of the BICAN atlas is focused on the basal ganglia and key associated areas^2,3^ based on sequencing over 17.4 million cells (Human 16.1M, Macaque 818K, Marmoset, 541K).

Advances in single-cell profiling technologies, neuronanatomy, and neuroinformatics have enabled development of two- and three-dimensional anatomic common coordinate frameworks (CCFs) for data reference and mapping^38–40^ (**Fig. 1D**.) The BICAN BG atlas is based on adult single-cell transcriptome-based cell classification and spatial mapping (**Fig.1E,F**), with each cell population additionally characterized using epigenomics^44^, chromatin conformation^45^ and spatial transcriptomics data^26,46^ with parallel sampling and integrated analysis across species. Multimodal characterization of electrophysiology and morphology using Patch-seq^6,47^ (**Fig. 1G**), in some cases leveraging cell type-specific viral genetic targeting, provides other essential cell type data for integration and comparison with molecular data. Data clustering and machine learning enable the construction of deep molecular taxonomies of cell types^48^ (**Fig. H**) that can be aligned across species with common ontology and nomenclature containing extensive cell phenotypic characterization with engaged community annotation. The atlas is mapped into species-specific anatomic frameworks with a novel cross-species unified and consistent structural ontology (**Fig. 1I**.) All data, tools, and supporting information are publicly available through the consortium portal (www.brain-bican.org), the Brain Knowledge Platform (BKP, knowledge.brain-map.org) data catalog^49,50^, and the BRAIN Initiative data archives (**Fig 1K**). A comprehensive list of BICAN resources is provided in **Suppl. Table 1.**

The BRAIN Initiative Cell Census Network (BICCN), the precursor to BICAN, achieved a comprehensive and high-resolution transcriptomic and epigenomic cell-type atlas for the adult brain of the mouse (*Mus musculus*), describing the transcriptomic and molecular signatures, spatial organization, and cell–cell interactions of thousands of cell types.^13,25,28^ Links between neuronal identity and axonal processes were characterized^27,51^ and the epigenetic landscape at regulatory elements and its connection with cell types were studied.^52,53,54^ Completing this work engaged a collaborative network of data-generating centers, data analysts, archives, and developers, with the goal of systematic multimodal brain cell type profiling and characterization^7^. However, the challenges of characterizing the cell-type structure of the mammalian brain are magnified substantially as brain size increases, introducing escalating conceptual, technical, and computational barriers. These difficulties are especially pronounced in the human brain, where several factors converge: (i) high biological complexity and inter-individual heterogeneity^55,56^, (ii) technical and experimental limitations related to tissue accessibility, postmortem quality, and donor variability^57,58^, (iii) conceptual and ontological challenges in defining cell types and constructing consistent, interoperable taxonomies and ontologies;^59^ and (iv) ethical and logistical considerations that constrain the scope and scale of human studies.^60,61^ Together, these issues create a level of complexity unprecedented in other mammalian systems and require coordinated conceptual, experimental, and computational innovation. The BICAN data ecosystem was developed as a multi-scale, multimodal, and multi-institutional effort to develop a standardized and reproducible framework to achieve the scientific goals of the consortium.

Standardization of protocols, metadata, and data-sharing practices has been central to genome science since the Human Genome Project and remains a defining feature of modern large-scale genomics efforts^62,63^. An important outcome of the present work is the development of analogous standards for all major attributes of cell type generation, classification, and annotation in the brain. These standards, ranging from coordinate systems through metadata and semantic frameworks to versioning and governance, parallel genomic standards now widely accepted in practice (**Box 1**). Each of these developed or adopted BICAN Cell Type Standards is discussed or referenced below.

### Box 1. BICAN Foundational Cell Type Standards.

Comparison of genomic standardization and cell type standardization frameworks. Abbreviations: CCF, Common Coordinate Framework; GIAB, Genome in a Bottle; GA4GH, Global Alliance for Genomics and Health; ABC Atlas, Allen Brain Cell Atlas; BKP, Brain Knowledge Platform; OBI, Ontology for Biomedical Investigations; EFO, Experimental Factor Ontology; HOMBA, Harmonized Ontology of Mammalian Brain Anatomy; INSDC, International Nucleotide Sequence Database Collaboration; HGNC, Human Genome Organization Gene Nomenclature Committee.

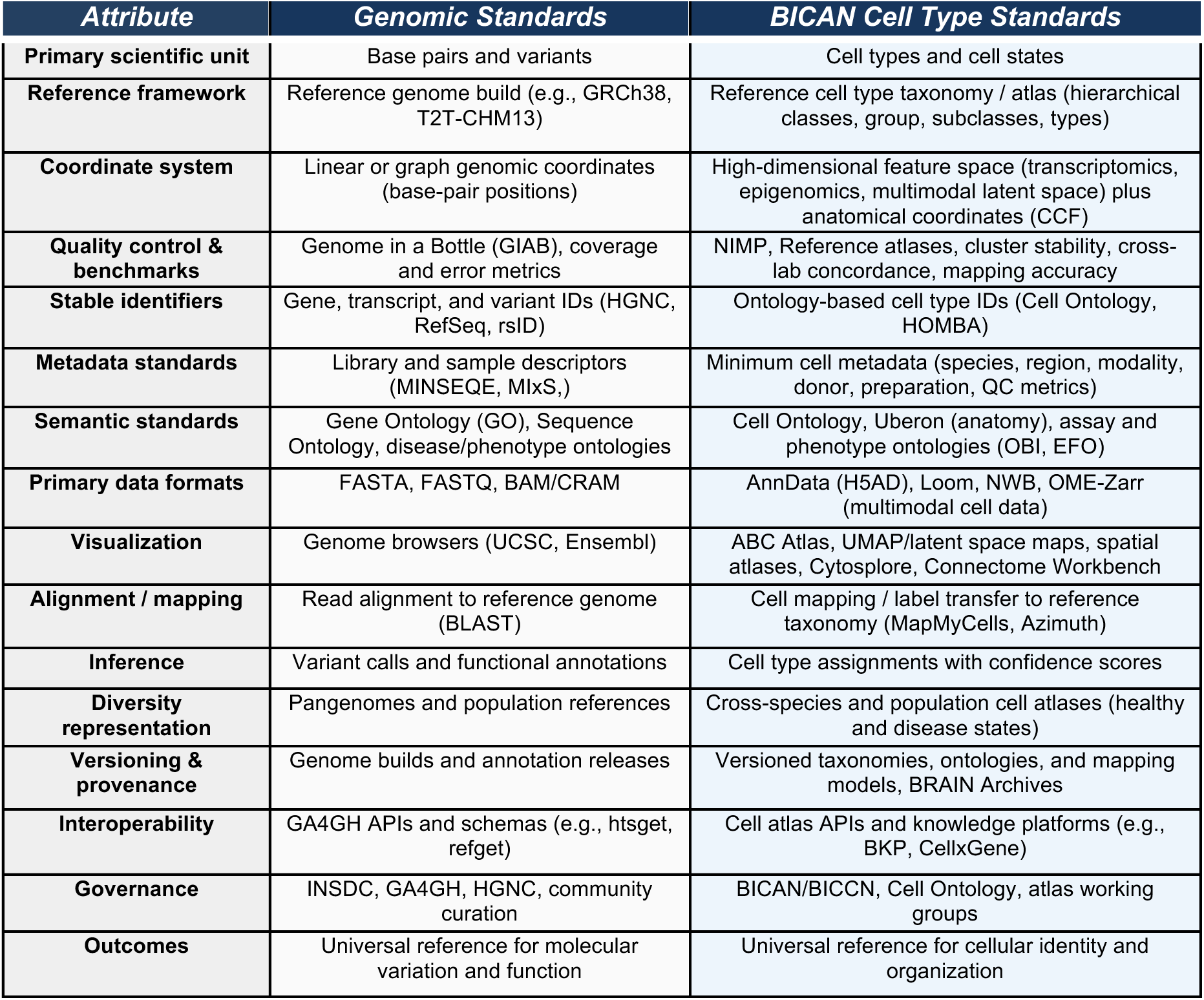

The BICAN ecosystem for generation, integration, and dissemination of brain cell atlases is a collaborative and actively managed multi-institutional effort (**Fig. 2**). Human and non-human primate brain tissue is acquired through coordinated donor programs and brain banks (including the NIH NeuroBioBank and UC Irvine’s BICAN Brain Donation Program) in a biospecimens core, with standardized protocols for tissue processing, preservation, and ethical/regulatory stewardship (**Fig. 2A**). These efforts ensure high-quality, well-annotated biospecimens suitable for downstream multimodal analyses. (see **Suppl. Information**). Biospecimens are processed within UM1 Centers(NIH Research with Complex Structure Cooperative Agreement) and U01 Collaboratories (NIH Research Cooperative Agreements) ^64^ as distributed research centers to generate high-resolution brain cell atlases (**Fig. 2B**). These efforts integrate molecular and anatomical signatures to define cell types, incorporating multispecies comparisons, developmental trajectories, and population-level variation. Standardized sequencing cores enable consistent production and harmonized metadata generation of large-scale single-cell and multiomic datasets across centers (**Fig. 2C**). Processed data through standardized mapping pipelines for each modality (Terra^65,66^) are deposited into coordinated public repositories, including NeMO (Neuroscience Multi-Omic Archive; single-cell omics), BIL (Brain Image Library; spatial transcriptomics and imaging), and DANDI (Distributed Archives for Neurophysiology Data Integration; electrophysiology, fMRI, and behavior), enabling broad access and interoperability (**Fig. 2D**). Central to the BICAN workflow are Coordinating Units for Biostatistics, Informatics, and Engagement (CUBIEs) that provide centralized infrastructure supports specimen tracking, sequencing workflow management, and data integration through interoperable portals, including specimen and sequencing management systems, as well as a brain knowledgebase for data access, FAIR tracking, annotation, mapping, and visualization (**Fig. 2E**).

**Figure 2.**
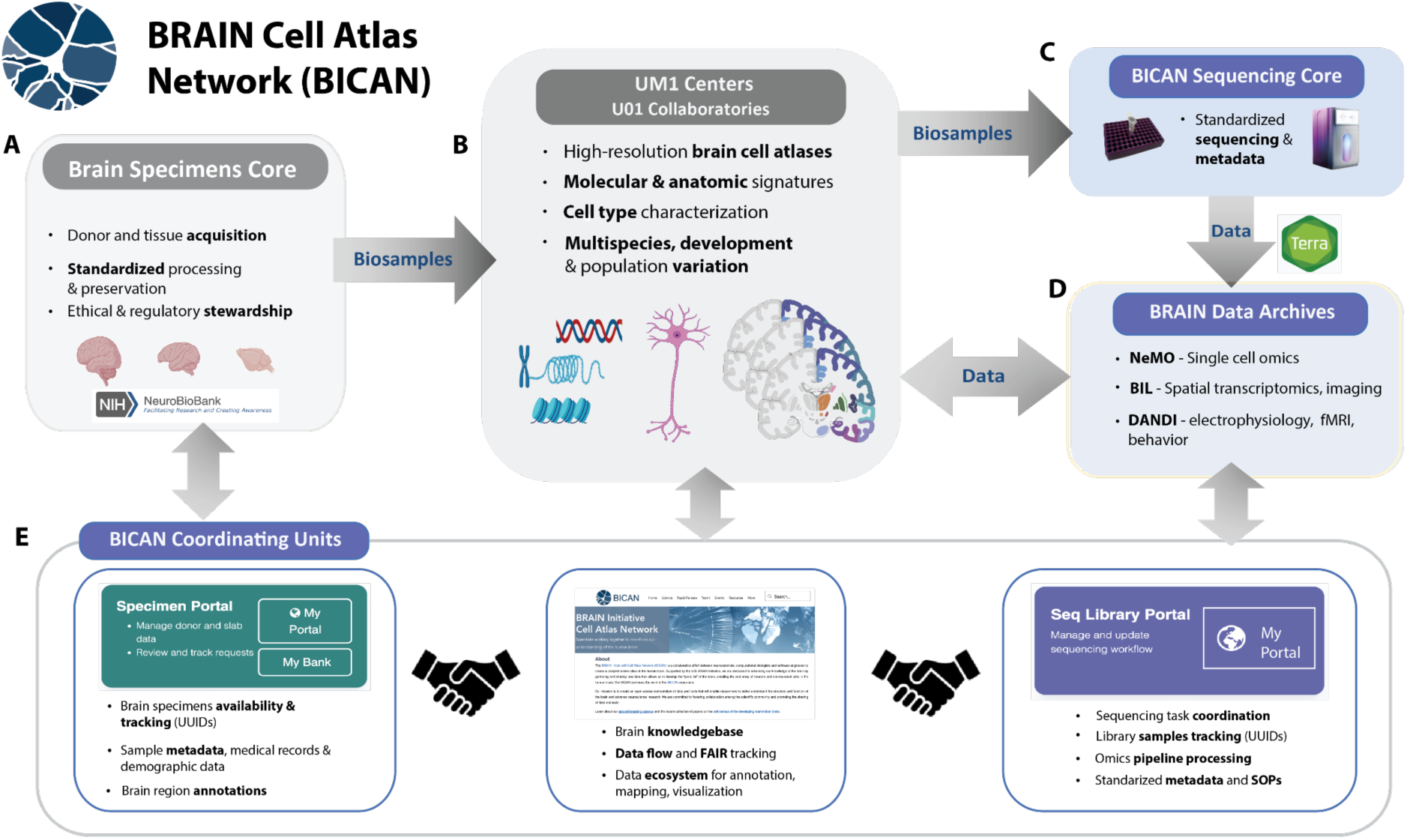
The BICAN ecosystem for generation, integration, and dissemination of brain cell atlases. The BICAN workflow engages A) Brain Specimens Core for tissue acquisition, neuropathology, and effective stewardship. B) UM1 Centers and U01 Collaboratories which generate data for molecular and anatomic signatures and cell type classification. C) Standardized sequencing used by UM1/U01 centers process data for genomic standardization (New York Genome Center^67^, Broad Institute Genomics Platform^65^. Sequence mapping through Terra^65,66^ produces standardized mapping and metadata. D) BRAIN data archives for single cell omics (NeMO), spatial transcriptomics and imaging (BIL), and electrophysiology and multimodal data (DANDI). E) BICAN Coordinating Units for Biostatistics, Informatics, and Engagement (CUBIE) that provide centralized infrastructure which supports specimen tracking, sequencing workflow management, and data integration^68^. Arrows indicate the flow of biospecimens and data through the system, from acquisition to atlas generation, standardized sequencing, and public data dissemination. Bidirectional connections reflect feedback between data generation, integration, and community use.

## 2. BICAN Basal Ganglia Atlases and Studies

BICAN UM1 centers (www.brain-bican.org) serve as the primary data generation and analysis units for this effort, producing comprehensive, high-resolution brain cell atlas datasets (**Fig. 3**). These coordinated activities span molecular, anatomical, and functional domains, enabling integrated characterization of the structure of the mammalian brain. Major initiatives include development of a functionally guided adult whole-brain cell atlas in human and non-human primates (Human and Mammalian Brain Atlas, Allen Institute)^69,70^ and large-scale multiomic profiling—including single-cell gene expression, chromatin accessibility, histone modifications, and DNA methylation—in human to define spatial organization and identify regulatory elements within non-coding DNA (Center for Multiomic Human Brain Cell Atlas, Salk Institute).^71–73^ The human brain exhibits substantial diversity in biological function and vulnerability to disease, yet the cellular and molecular basis of inter-individual variation remains poorly understood.^74,75^ The Atlas of Human Brain Variation (Broad Institute)^76,77–80^ addresses this gap by applying single-cell and spatial genomics across hundreds of individuals to distinguish stable versus dynamic molecular and spatial features of cell types and to quantify the influence of human genetic variation on cellular phenotypes.

**Figure 3.**
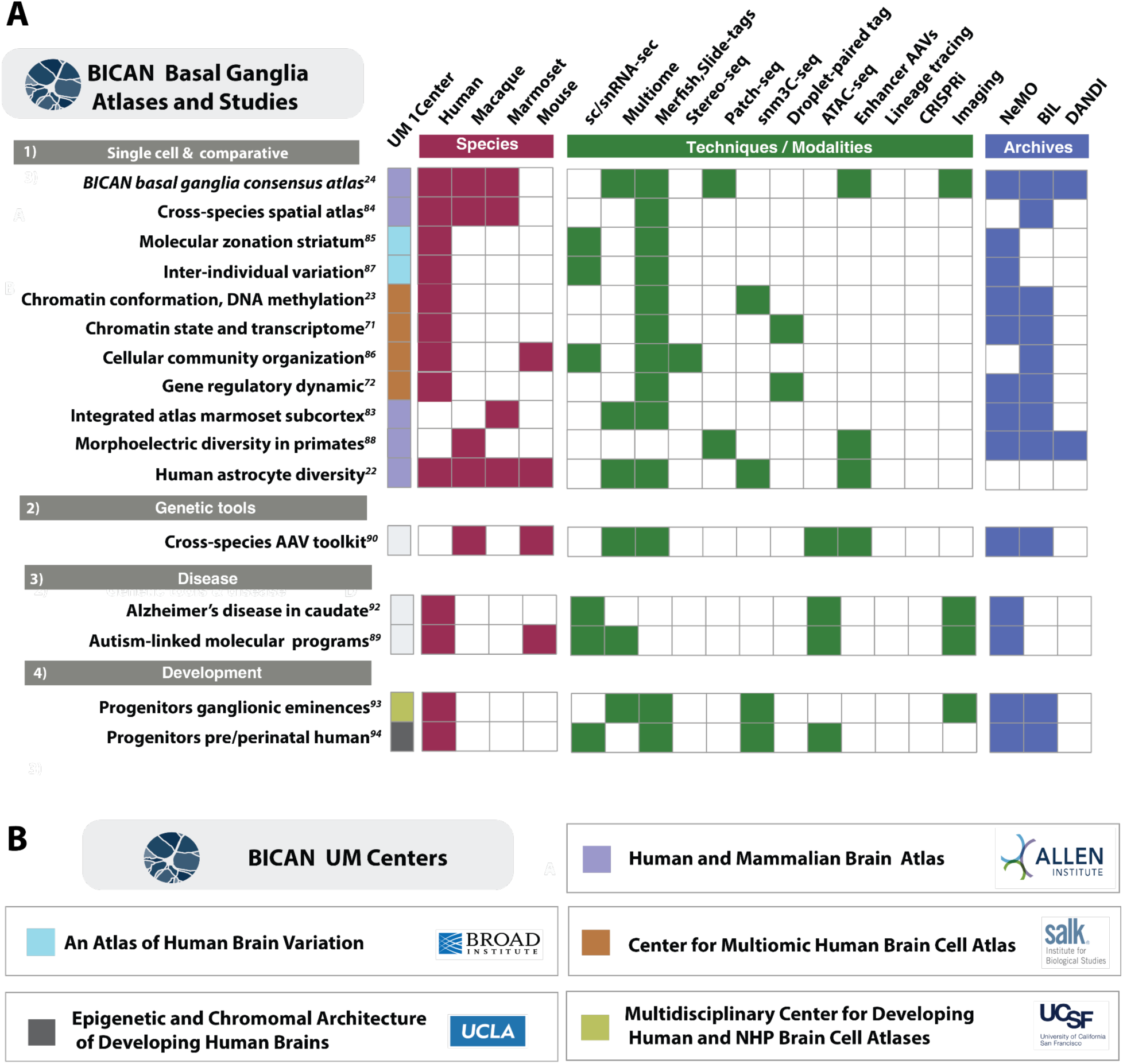
BICAN Basal Ganglia Atlases and Studies. A) BICAN Basal Ganglia Atlases and Studies. Studies referenced in text grouped by 1) primary atlas studies, 2) multimodal cellular properties and comparative analyses, 3) genetic tools and disease applications and 4) developmental studies. For each study, the matrix shows the species profiled, primary modalities and techniques used, and the BRAIN Initiative data archive where primary data is found (NeMO: Neuroscience Multi-Omic Archive, BIL: Brain Image Library, DANDI: Distributed Archives for Neurophysiology Data Integration.) B) BICAN UM Centers (www.brain-bican.org). The BICAN consortium is represented by several centers for single-cell transcriptomic, epigenomic, and multimodal data profiling. The five UM centers contributed to the basal ganglia studies described in A) with specific studies identified by color code.

Developmental processes further contribute to cellular diversity. Cortical neuron identities are specified through hierarchical programs involving lineage, temporal progression, and transcriptional regulation^81^. While adult cell types are increasingly well characterized, transitional cell types and dynamic states during human brain development remain incompletely defined. BICAN is profiling anatomically distinct regions of developing human, rhesus macaque, marmoset, and mouse brains across key developmental epochs, including mid-gestation, neonatal, childhood, and adolescence. The Multidisciplinary Center for Developing Human and NHP Brain Cell Atlases (University of California, San Francisco; USCF)^77–79^ focuses on identifying transient cell populations and reconstructing developmental trajectories, while the Center for Epigenetic and Chromosomal Architecture of Developing Brains (University of California, Los Angeles; UCLA)^77,78^ integrates epigenomic and three-dimensional chromatin architecture to define regulatory programs underlying lineage commitment and maturation. The Center for Comprehensive Single-cell Atlas of the Developing Mouse Brain (Harvard/Allen/UCSF) is creating a multimodal developing cell atlas for the whole mouse brain at all pre- and postnatal timepoints.^49,50,82^ Together, these coordinated efforts generate and integrate multimodal datasets across species, developmental stages, and individuals to identify conserved and divergent features of brain organization and define cell types and regulatory mechanisms relevant to both healthy function and disease.

Through these studies, BICAN integrates a diverse set of data-profiling and informatics initiatives spanning single-cell and spatial genomics analyses of the basal ganglia^24,83^. The studies summarized in **Fig. 3A** are organized by species (red), experimental modality (green), and data archive location (blue). Together, these provide a comprehensive molecular and cellular characterization of the primate basal ganglia and related brain structures. By integrating spatial transcriptomics, single-cell multi-omics, epigenomics, electrophysiology, and cross-species comparative analyses, this body of work establishes a unified framework linking cell identity, spatial organization, gene regulation, development, evolution, and disease vulnerability across the primate brain.

Joint accomplishments of this work include a complete aligned multispecies taxonomy of the mammalian basal ganglia^24^ characterizing homologous and species-specific cell types. Several studies aligned molecular data to this reference (**Fig. 3A-1**) using single-cell and spatial transcriptomic analyses^84^ to reveal previously unrecognized mesoscale organization within basal ganglia circuits, including spatial gradients, striosome–matrix compartmentalization^85^, molecular zonation of the human striatum across cell types and people^85^, and regionally specialized cellular communities that link molecular identity with anatomical and circuit context^86^. A study on inter-individual variation of cellular properties of the human striatum identified thousands of age-associated (but few sex-associated) variations in gene expression^87^. Many of these effects of age were cell-type-specific, and individuals’ ages could be predicted to within about five years based on RNA-expression patterns from any of the striatal cell types.

Complementary epigenetic studies with mapped data extend these atlases into the regulatory domain by jointly profiling chromatin accessibility, histone modifications, DNA methylation, three-dimensional genome organization, and gene expression.^23,71^ These data identify cell-type-specific regulatory programs, enhancer–promoter interactions, and transcription factor grammars underlying neuronal and glial identity. The resulting regulatory architecture combines conserved transcriptional motifs with region-specific regulatory elements and developmental state transitions^72^. Multimodal approaches further refine and validate neuronal classification where techniques such as Patch-seq integrate morphology, electrophysiology, and transcriptomics from individual neurons, demonstrating morphoelectric properties, including dendritic architecture and intrinsic excitability that align with transcriptomic subclasses while contributing additional axes of diversity^88^.

Comparative analyses across human, macaque, marmoset, and mouse datasets revealed both conserved and species-specific cellular and regulatory features^24,83^. Major neuronal classes, including medium spiny neurons, are evolutionarily conserved, whereas primates exhibit expansions and specialization of neuronal and interneuron populations. Cross-species regulatory analyses further identify conserved enhancer grammars associated with cell identity while highlighting evolutionary modifications of transcriptional networks^24,83^and evidence that human basal ganglia astrocyte diversity reflects evolutionary and circuit-level specialization^84^.

The use of snATAC-seq and 10x multiomic datasets for the discovery of basal ganglia cell type enhancers led to a cross-species enhancer-based adeno-associated virus (AAV) toolkit enabling cell-type-specific targeting across basal ganglia circuits (**Fig. 3A-2**).^88–90^ Studies with a disease-focused analysis (**Fig. 3A-3**) include a cellular-resolution atlas of Alzheimer’s disease–associated pathology in the caudate nucleus was generated, leveraging the new basal ganglia cellular taxonomy^92^, and a characterization of autism-associated disruption of cell-type-specific molecular programs in the putamen.^89^

Developmental studies **(Fig. 3A-4**) examined the ontogeny of basal ganglia cell types. These include analyses of progenitor diversity in the developing human ganglionic eminences^93^; multi-omic characterization of transitions in striatal and cortical progenitors in the prenatal and perinatal human brain, barcoded lineage tracing of interneuron fate specification during primate basal ganglia development, and a single-cell epigenomic and chromatin-architecture atlas of developing basal ganglia and inhibitory neurons.^91,94^ The breadth of these studies provides the data and tools for demonstrating a comprehensive analysis of human and non-human primate basal ganglia and resources for further study of this key brain system.

**Figure 4.**
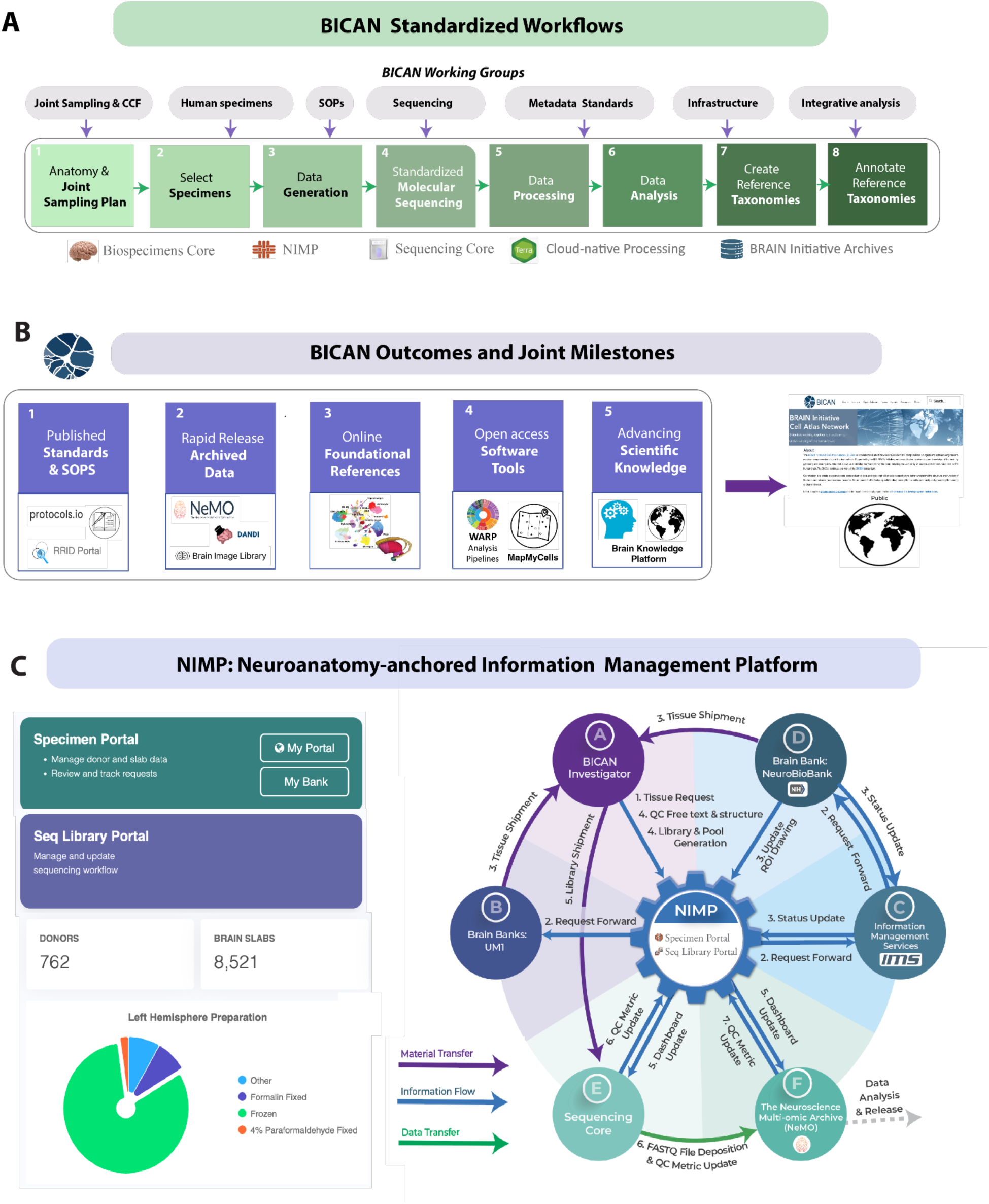
BICAN Standardized Workflows and Outcomes. A) The BICAN ecosystem operates to turn the consortium’s research into a coherent and scalable system based on standardized workflows and coordinated joint milestones and outcomes: A1) development of a consistent cross-species joint sampling plan, A2) specimen selection, (A3-A8) data generation through annotated reference taxonomies. Cross-institution working groups develop protocols and standards for joint sampling, specimen selection, standard operating procedures (SOPs), metadata standards, software infrastructure, and integrative analysis. B) BICAN consortium outcomes are coordinated through joint milestones that represent major deliverables. C) The Neuroanatomy-anchored Information Management Platform (NIMP^68^, RRID:SCR_024682), a web-based application for managing the integrative workflows from BICAN, is a cornerstone of the BICAN ecosystem. D) NIMP manages initial operations of BICAN workflow (A1-A4) from the selection of human donors from the NIH Neuro Biobank, tracking sequencing status from BICAN-designated sequencing centers, and porting study metadata for downstream data releases. A public version of NIMP is accessible through NIMP Analytics (SCR_028218) without a user account.

## 3. The BICAN Cell Atlas Ecosystem

Achieving multiscale and multimodal profiling of the human brain further requires cross-disciplinary coordination and infrastructure capable of integrating diverse data types into coherent, reproducible frameworks^7^. The BICAN ecosystem operationalizes this integration through standardized workflows, coordinated milestones, and shared analytical frameworks (**Fig. 4A**). The overall workflow begins with a consortium-wide anatomical joint sampling strategy, followed by biobank specimen acquisition and preparation^95^, data generation across participating centers, and standardized molecular profiling through centralized sequencing platforms, including the New York Genome Center^67,96^ and the Broad Institute Genomics Platform^67,96^.

Subsequent standardized data processing, integration, and analysis support the construction of cell type taxonomies and the generation of annotated reference atlases, which are released as accessible data products (**Fig. 4A**,1-8). This process is guided by consortium working groups that actively coordinate specimen selection, metadata harmonization, infrastructure development, and integrative analysis. Collectively, these efforts deliver a set of coordinated outcomes (**Fig. 4B)**, including: (i) standards and standard operating procedures; (ii) rapid and open data release; (iii) foundational reference resources, including common coordinate frameworks (CCFs) and ontologies; (iv) open-access software tools; and (v) an integrated platform for knowledge organization, access, and visualization. Together, these components constitute the publicly accessible outputs of BICAN and establish a scalable foundation for atlas-driven neuroscience.

### Human Brain Specimens for BICAN

Access to diverse, high-quality human brain specimens is essential for the success of the BICAN program^95^. Partnerships with brain banks, advances in tissue collection, and quantitative neuropathology enable access to well-characterized donors and precisely annotated specimens. Achieving BICAN objectives requires preservation methods compatible with molecular assays, as well as harmonized procurement protocols and donor metadata standards(see Metadata-schemas^97^) to support mapping into common coordinate frameworks. BICAN neuropathology teams optimize tissue processing to balance broad coverage of major cell types with deep characterization of rare populations, enabling high-resolution, anatomically grounded cell atlases (**Suppl. Information**).

To meet these requirements, BICAN established a modernized framework for human brain donation that integrates coordinated donor networks, standardized consent and de-identification for open-access genomic data, and validated protocols for tissue procurement, dissection, and preservation across multiple tissue banks. Three-dimensional brain reconstructions and rapid, portable MRI are used to align donor brains to common coordinate frameworks, improving anatomical precision and guiding tissue dissection (**Suppl. Information**). A standardized donor characterization pipeline harmonizes minimum data elements across tissue banks, integrating clinical and neuropsychological history, exposure and toxicology data, infectious disease screening, neuroimaging, and comprehensive neuropathological evaluation. The pipeline also addresses ethical considerations, including potential biases in donor characterization and allocation.

In this context, the specimen pipeline functions as the foundational layer of the atlas, ensuring that biological variability is captured in a standardized and spatially grounded manner suitable for integration into a reference framework. Together, this establishes a scalable, high-quality biorepository that supports the generation of BICAN reference atlases and provides a foundation for studies of the human brain.

### Neuroethics Engagement

The BICAN program’s specimen collection, analysis, and data curation pipeline comes with ethical and social challenges for engaging respectfully with human donors and their communities, collecting heterogeneous cohorts of donors, and ensuring broader access to software tools and archived data. To address these challenges, bioethicists were integrated into the human tissue procurement pipeline, leadership activities, and the broader BICAN team. The neuroethics team leads engagement efforts through several activities, including ethics engagement surveys, advisory board meetings, reading groups, topical task forces, public outreach events, and 1-on-1 consultations. This approach has fostered a culture of ethics engagement across the program.

### NIMP - Neuroanatomy-anchored Information Management Platform

A cornerstone of the BICAN ecosystem is the Neuroanatomy-anchored Information Management Platform^68^ (NIMP), a web-based digital twin platform for managing BICAN brain specimen selection and library processing. The NIMP is an online cross-consortium laboratory management system for BICAN brain specimen selection and library processing and is essential to BICAN’s operational workflow. NIMP consists of two portals for BICAN collaborative data generation: the Specimen Portal and the Sequence Library (SeqLib) Portal (**Fig. 4C)**. The Specimen Portal focuses on tissue management from donors to brain slabs and annotated brain samples. The SeqLib Portal manages the workflow starting from tissue, and downstream to track data deposition to assay-dependent, data-modality-specific archives.

The NIMP portals coordinate the full BICAN processing workflow from relationships with NeuroBio Bank, specimen selection, and annotation and coordination where APIs allow seamless interfacing with upstream human donor resources from the NIHNeuro Biobank, tracking sequencing status from BICAN-designated sequencing centers, and porting study metadata for downstream data releases through the Neuroscience Multi-omic (NeMO) Data archive (RRID:SCR_016152) (**Fig. 4D**). To ensure data FAIR-ness, NIMP operates an interactive provenance-graph visualization engine, allowing step-by-step inspection of the end-to-end data generation process using Sankey diagrams ^98^ that represent distinct types of intermediate resources generated in the process. The NIMP technique to achieve this donor-to-data, end-to-end provenance tracking is through the use of NHash Identifier (RRID:SCR_025313) which operates a simple block-chain strategy^68^ encoding the direct dependencies among the resource types.

### Uniform and Centralized Molecular Data Processing

BICAN uses standardized molecular sequencing across all projects through the Broad Institute Genomics Services ^99,100^ and New York Genome Center^67^. After prepared library aliquots are sequenced, molecular data is immediately transferred to the NeMO Archive. Raw data is first checked for quality assurance and checking for expected descriptive metadata, file types, and checksums. These raw sequencing data are made available to the generating labs for any additional inspection. Data that passes NeMO quality checks are then processed using horizontally scalable, cloud-native pipelines using Broad Terra (RRID:SCR_021648; terra.bio). To establish scalable data operations, the NeMO Archive (RRID:SCR_016152) employs federated integration with Terra where pipelines are executed on a commercial cloud platform (Google Cloud Platform^101^). These data operations are scripted to enable processing high volumes of data. Integration between the NeMO archive (where data is stored) and Terra (where data is processed) enables efficient data processing without movement of the original raw data. After processing is complete, the resulting analysis data is moved to NeMO where it is once again checked for quality assurance. Consortium-defined uniform processing pipelines consistently generate over 100 standard quality metrics focused on the quality of libraries, cells, and genes. A subset is extracted from the derived data and stored at NeMO and NIMP for investigators to confirm data quality. Once confirmed, those uniform data are flagged for the release process.

BICAN has adopted and improved scalable technology platforms and multiplex assay protocols to achieve high production and cost efficiency, enabling the consortium to generate large-scale molecular and anatomical data that meets Complete, Accurate, and Permanent (CAP)^74^ criteria. BICAN uniform molecular data processing pipelines are developed by the Broad Institute in close collaboration with BICAN scientists across institutes. These consortium data processing pipelines (WARP^102^) include pipelines for processing 10X Multiome (3’ transcriptomics and ATAC-seq; RRID:SCR_024217, Paired-Tag 3’ transcriptomics and histone modification RRID:SCR_025042, snm3C-seq (methylome and chromatin contact; RRID:SCR_025041), Smart-seq (full transcript, Patch-seq; RRID:RRID:SCR_018920), and Slide-seq/Slide-tag (spatial transcriptomics; RRID:SCR_023379/SCR_027567) data as well as standard reference preparation (BuildIndices; RRID:SCR_027630). Working groups for each pipeline and core reference (human, mouse, marmoset, and macaque) were established (**Fig. 4A**). Processing pipelines were defined and optimized (for science, speed, and cost) by Broad pipeline engineers working with consortium scientists focused on defining and validating the scientific rigor of the produced data.

## 4. BICAN Basal Ganglia Ontology and Coordinate Frameworks

The basal ganglia are a group of deeply located subcortical nuclei that form recurrent cortico–basal ganglia–thalamo–cortical loops and play central roles in motor control, action selection, procedural learning, reward processing, and cognition^103,104^. The composition of the basal ganglia varies across anatomical, functional, and clinical frameworks, reflecting differences in historical and experimental context ^105,106^. To emphasize connection with other major consortia (e.g., BRAIN CONNECTS), BICAN enhances understanding of the classical striatal-pallidal complex (caudate nucleus, putamen, and globus pallidus) with an emphasis on circuit-based structures (subthalamic nucleus, substantia nigra), and related ventral tegmental area, as these midbrain and diencephalic structures are indispensable components of canonical basal ganglia loops and intrinsic and output nodes (**Fig. 5A,B)**.

**Figure 5.**
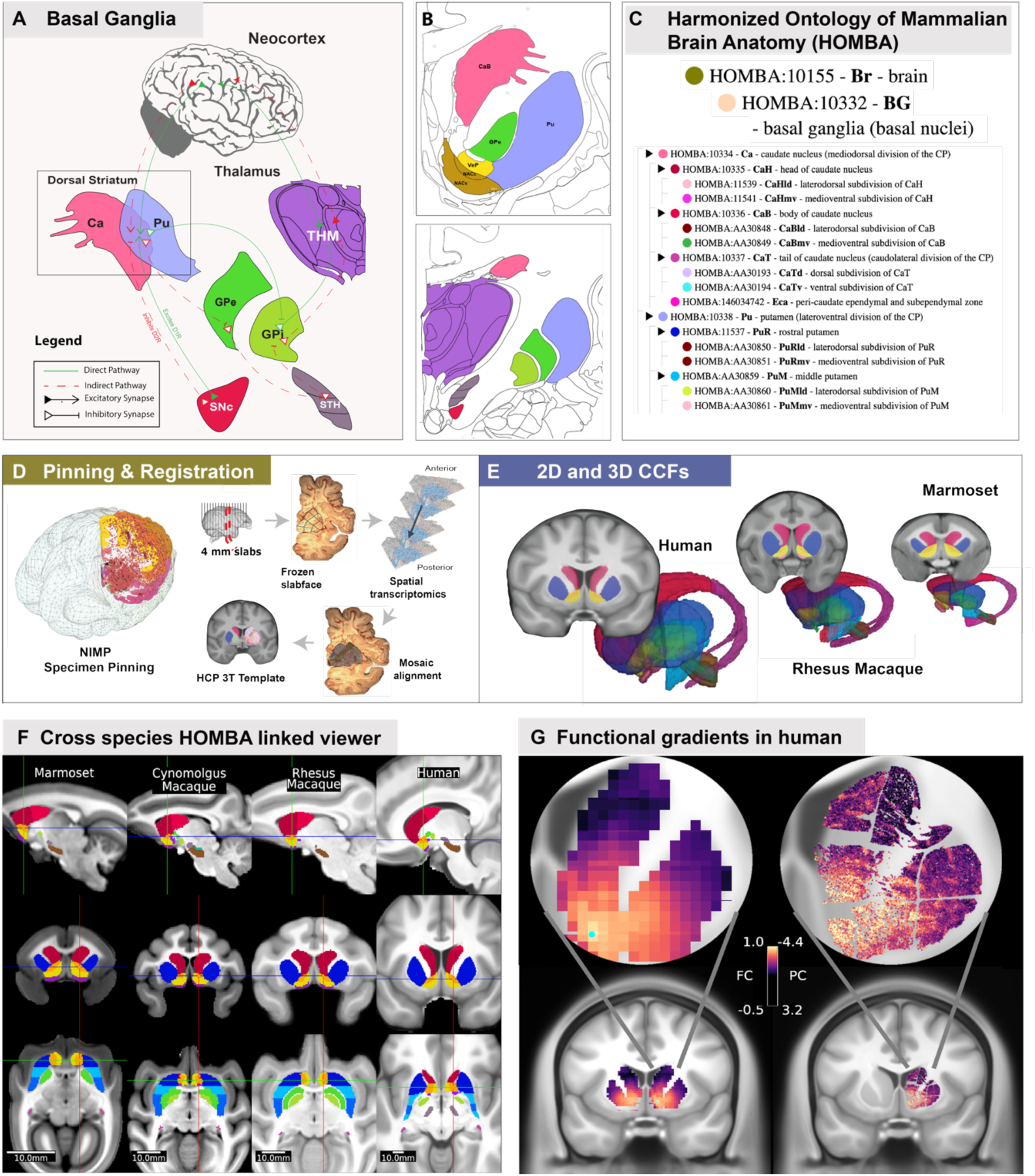
Basal ganglia anatomy, HOMBA, CCFs, and spatial mapping. A) BICAN enhances understanding of the classical striatal-pallidal complex (caudate nucleus, putamen, and globus pallidus) with an emphasis on circuit-based structures. B) Annotated plates from the Allen Human Reference Atlas^107^ showing basal ganglia structures. C) Harmonized Ontology of Mammalian Brain Anatomy (HOMBA), a cross-species whole brain ontology. D) Tissue sampling and registration workflow for basal ganglia spatial transcriptomics atlas. Profiled tissue samples are pinned based on HOMBA ontology. For spatial data the brain is partitioned into 4mm-thick frozen slabs, which are subdivided into ∼1cm² blocks along anatomical boundaries to accommodate MERSCOPE imaging constraints. Blocks are sectioned and processed for spatial transcriptomics. Following imaging, spatial transcriptomics data from each section are registered to block-face images, then aligned and stitched back to the corresponding slab-face image. The reconstructed sections are then registered to the HCP 3T template which has been registered into the MNI152 Non-linear 6A Asymmetric coordinate space and annotated with basal ganglia structures using HOMBA nomenclature (**Suppl. Information**). E) Structures shown for basal ganglia.

### Harmonized Ontology of Mammalian Brain Anatomy (HOMBA)

A novel ontology integrating human and non-human primate anatomy through cross-species common coordinate frameworks was developed using unified structural ontologies and nomenclature (HOMBA, **Fig. 5C**). HOMBA is created using multiple/multimodal datasets across species, in which the ontology respects similar cyto- and chemo-architecture across species, and conserved gene markers (RRID:SCR_027628) where homologous structures are labeled with the same anatomical ontology/terminology. BICAN’s joint sampling plan is based on the HOMBA ontology to generate the data needed to produce a comprehensive cytoarchitectural map of the human and NHP brain. Key points included: (i) generating a consistent anatomical ontology for sample dissections, to ensure downstream data can be harmonized across data profiling UM groups, (ii) removing redundancy between independently generated plans, and (iii) identifying opportunities to create more synergies between the independent group plans.

### Common Coordinate Frameworks in BICAN

A central priority of BICAN is to localize transcriptomic tissue samples to well-defined, functionally relevant brain structures with the highest possible accuracy, subject to the practical constraints and limitations inherent to individual datasets. Integration with other datasets of the same and different data modalities is enabled by effective imaging data analysis pipelines which process and rapidly share verified raw and derived data, perform data analysis, visualization, and mapping to common coordinate systems. For the BICAN basal ganglia consensus atlas, a population-average template is generated for each species **(Fig. 5D**) using structural MRI scans from multiple individuals registered into a common spatial framework. Each common coordinate framework (CCF) necessarily incorporates two complementary coordinate systems: a three-dimensional volumetric coordinate system optimized for representing subcortical domains with high anatomical fidelity, and a surface-based coordinate system optimized for representing the cerebral cortex, which is a sheet-like structure that is highly convoluted and, in humans, exhibits substantial inter-individual variability^110^.

Cross-species atlases for human, rhesus macaque, cynomolgus macaque, and common marmoset were constructed from MRI-based anatomical templates. The human atlas used the recently released Human Connectome Project Young Adult (HCP-YA 2025) template, which is a group-averaged 3T MR of 1071 subjects, aged 22-35. This template was registered to the MNI152 6th generation non-linear asymmetric space. Macaque and marmoset MRI templates were generated specifically for this study and were based on cohorts of 25 individuals per species^116^.Each MRI template was parcellated using HOMBA nomenclature. Atlas data assets were organized according to the AtOM data model^117^, which conceptualizes an atlas as an integrated set comprising an anatomical reference template, a coordinate space defining mathematical and anatomical axes, an annotation set that parcellates the template into regions, and a terminology that assigns human-readable labels and associated metadata. Within BICAN, templates, coordinate spaces, and annotation sets are species-specific, whereas terminology is shared across species. This atomic and modular data model facilitates interoperability, extensibility, and version control.

All atlases are packaged in cloud-native streaming formats and made available (**Suppl. Resource Table)** for interactive visualization using Neuroglancer^118^, which supports image volumes, three-dimensional meshes, and point-cloud annotations. Existing subcortical annotations for marmoset, macaque, and human are distributed across disparate reference images, frequently employ inconsistent nomenclature for homologous structures, and often subdivide structures at different levels of granularity, thereby complicating cross-species comparisons. To address these limitations, annotations were transferred onto state-of-the-art group-average reference images, harmonized structure names, and embedded all nomenclatures within a unified cross-species hierarchical taxonomy. Visualization using Connectome Workbench^111^ enables intuitive navigation across species **(Fig. 5F**) and supports robust indexing across differing annotation granularities by mapping selected substructures (e.g., head of the caudate nucleus) to their homologous parent structures (e.g., caudate nucleus) when necessary.

Histologically informed 3D annotations parcellate the MRI templates in each species into regions named by HOMBA, which harmonizes labels and colors across species. Parcellations are shown as 3D volumes and meshes and can be viewed interactively in Neuroglancer^108^ ^109^. F) Point and click cross-species navigation in Connectome Workbench^111^ of four hierarchical basal ganglia volume annotations in common marmoset, cynomolgus macaque, rhesus macaque, and human. Group average T1w structural MRI underlays. G) Functional imaging and spatial transcriptomics. Comparison of MRI and STx ^84^ registered to a common coordinate framework. Left, group average human 7T rs-fMRI functional correlation (FC) from seed in ventromedial striatum, only subcortical gray matter voxels are shown. Right, dots representing cells in human donor basal ganglia colored by gene expression principal component (PC) score. Group average 3T T1w structural MRI underlays.

### *In vivo* functional and *ex vivo* transcriptomic gradients

Mapping multimodal data to CCF space permits direct comparisons of *in vivo* functional and architectonic MRI features and *ex vivo* transcriptomic features. Through the alignment and registration of spatial transcriptomic data derived from slabfaces and registered to the HCP human template with HOMBA annotations (**Fig. 5E, Suppl. Information**) functional and transcriptomic structure can be compared. Functional dorsolateral to ventromedial gradients in the striatum have been shown to correspond to motor to associative and limbic function^119,120^. The BICAN cross-species spatial atlas identifies transcriptomic gradients in human basal ganglia superimposed on discrete compartments^121^. To illustrate BICAN atlasing standards, **Fig. 5G** shows a qualitative correspondence in a ventromedial-dorsolateral gradient observed in (left) group average human 7T resting state MRI functional connectivity from a seed in ventromedial striatum and (right) a principal component of spatial transcriptomic-derived gene expression in a human donor^84^. Data have been co-localized to the same oblique near-coronal slice in CCF space and overlaid on top of a group average 3T T1w structural MRI reference image.

## 5. BICAN Basal Ganglia Taxonomy

Advances in high-throughput transcriptomic profiling and machine learning have transformed our capacity to systematically classify these neuronal populations and integrate taxonomies across species. With roughly 200 million neurons in the human BG alone, these nuclei exhibit remarkable cellular and molecular diversity that has historically been defined through cytoarchitectural, connectivity, neurophysiological, and molecular criteria. Building on this foundation, a unified, cross-species taxonomy of the mammalian basal ganglia was developed through coordinated efforts of the NIH BRAIN Initiative, BICAN Human and Mammalian Brain Atlas (HMBA)^70^ and the Armamentarium Basal Ganglia AAV Toolbox. ^70,122^

The *HMBA cross-species consensus taxonomy of basal ganglia* (**Figure 6)**^123–125^ integrates established nomenclature from prior literature with new single-nucleus RNA-sequencing data from human, macaque, marmoset, and mouse, yielding a standardized framework grounded in conserved molecular signatures and evolutionary relationships. By harmonizing terminology, marker gene definitions, and hierarchical cell-type relationships across species, the HMBA consensus taxonomy enables consistent communication about cell types, facilitates the development of genetic tools for selective cell targeting and perturbation, and provides a foundational resource for studying the organization and function of the basal ganglia across mammals.

**Figure 6.**
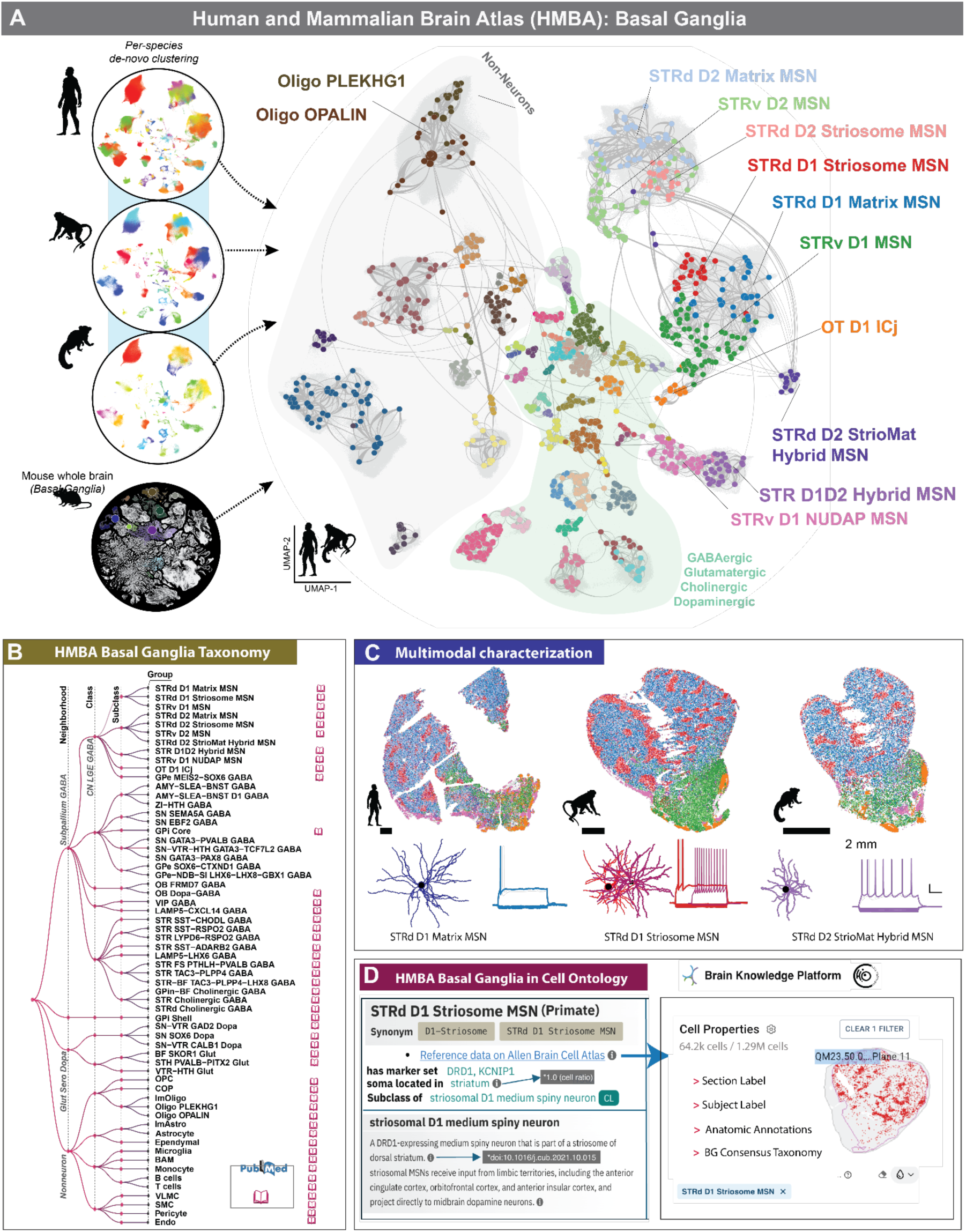
BICAN Basal Ganglia Molecular Reference Standard. A) The BICAN consensus basal ganglia cell type taxonomy^24^ generated through iterative clustering and cross-species integration of single-nucleus transcriptomic data from human, macaque, and marmoset spanning key HOMBA structures including the caudate (Ca), putamen (Pu), nucleus accumbens (NAc), external and internal globus pallidus (GPe, GPi), ventral pallidus (VeP), subthalamic nucleus (STH), and substantia nigra (SN). Combined with whole-brain transcriptomic datasets from the mouse, these data support a unified taxonomy that resolves conserved cell types while revealing species-specific molecular divergence. B) Cell type definitions are validated and refined through integration with additional datasets with a standardized nomenclature. C) Multispecies spatial transcriptomics, along with Patch-seq-based morphology and electrophysiology in macaque, are used for validation and refinement of the taxonomy. D) Integration with the Cell Ontology supports classification describing the general properties of cell types using data from multiple references and covering multiple modalities.

To support this taxonomy, the team compiled comprehensive metadata describing each identified cell type, including gene-expression profiles, marker genes derived from snRNA-seq analyses, and synonymous nomenclature drawn from previous studies. ^13,55^ These metadata are organized under an aligned Allen Institute Taxonomy (AIT)^126,127^ data schema, ensuring compatibility with community platforms such as CELLxGENE and the Cell Annotation Platform (CAP)^128^, and enabling seamless integration and reuse across ongoing BICAN efforts.

### Clustering and cell type annotation across species

The BICAN consensus basal ganglia cell type taxonomy was generated through iterative clustering and cross-species integration of single-nucleus transcriptomic data obtained using the 10x Genomics Multiome platform^129^. The taxonomy incorporates snRNA-seq data from human, macaque, and marmoset spanning key HOMBA structures including the caudate (Ca), putamen (Pu), nucleus accumbens (NAc), external and internal globus pallidus (GPe, GPi), ventral pallidus (VeP), subthalamic nucleus (STN), and substantia nigra (SN). Combined with whole-brain transcriptomic datasets from the mouse^13,28^, these data support a unified taxonomy that resolves conserved cell types while revealing species-specific molecular divergence (**Fig 6A**). The resulting hierarchy was validated through marker gene expression analysis, comparison with previously published taxonomies, and self-projection across species, ensuring both the biological accuracy and robustness of each level within the taxonomic framework ^130^.

A standardized pipeline for taxonomy development was established and adopted across BICAN groups involved in developing the consensus taxonomy to generate harmonized human, macaque, and marmoset cell type taxonomies.^131^ Building on the robust clustering workflow used in the mouse whole-brain atlas study^13,130^, the pipeline was extended to primates by incorporating scVI models^132^ to correct for donor-specific variation, a critical confounding factor in human and NHP datasets. Single-nucleus RNA-sequencing data from each species were clustered and low-quality nuclei were removed. Cluster annotations were guided by existing taxonomies and references.^13,55,133,134^ These initial annotations provided a foundation for cross-species mapping and nomenclature standardization. While each species was annotated independently, the taxonomies were reciprocally mapped and aligned across the species to ensure consistency and establish cell type homologies. Additional literature mining identified canonical basal ganglia cell types that were underrepresented or absent from single-cell atlases, which were adopted in the BICAN consensus taxonomy.

Following the generation of this initial annotated taxonomy, cell type definitions were validated and refined through integration with additional datasets, including spatial transcriptomics, Patch-seq, and population-scale snRNA-seq on >150 donors. At this stage, the taxonomy was iteratively refined by manual curation among collaborating teams and computational reassessment (**Fig. 6B**), a process essential to reconcile transcriptomic and anatomical definitions. Spatial transcriptomics identified anatomically organized cell populations and hybrid cell types (**Fig. 6C**) such as the striosome and matrix medium spiny neurons (MSNs). A more detailed consortium analysis performed with Slide-tags revealed a conserved, broader zonation pattern observed in each subset of MSN.^85^ Three cholinergic RNA-seq clusters were further resolved into spatially distinct subtypes, and multiomic signatures combined with anatomical localization confirmed that *GPi Core* and *GPi Shell* cell types are spatially distinct and conserved cell types across species.

Electrophysiological measurements using Patch-seq experiments revealed electrophysiological and morphological distinctions between matrix, striosome, and hybrid MSN populations (**Fig. 6C**). Applying label transfer to a population-scale dataset of more than 150 donors confirmed the consistency and conservation of the taxonomy in a broader postmortem sample, and identified patterns of cell-type-specific variation across people.^87^ Together, these efforts established a reproducible framework for cross-species refined taxonomy combining computational integration that anchors non-omics modalities and achieves a more complete and functionally grounded consensus taxonomy of the basal ganglia (**Suppl. Information**).

### BICAN Taxonomies and Cell Ontology

Brain cell type taxonomies are a core product of BICAN. They provide standard names and hierarchy for cell types linked to reference data and analysis across species and modalities. Annotation transfer tools such as Azimuth^135^ or MapMyCells^136^ support transfer of these names between datasets. However, on their own, they are not suitable for use by platforms and tools that rely on standard ontologies for integration of cell types across all data sources and species. For example, the CELLxGENE Discover, CellGuide platforms, and Census API^137^ use the Cell Type Ontology to standardize cell type annotation and to drive search, browsing, API queries, and data aggregation for marker calling. The Cell Type Ontology (CL;RRID:SCR_004251)^138^ provides a structured and standardized vocabulary for representing cell types across species and is used by a wide range of platforms and tools to provide standardized cell type annotation. CL is also an integral part of the Human Cell Atlas (HCA) integrated atlas efforts. It provides a standard for annotating transcriptomic types in HCA and other atlases, and the atlases in turn provide new terms for CL.

To support broader community use of BICAN cell typing standards and integration with other standard atlasing efforts, all cell types defined by BICAN taxonomies are being integrated into the Cell Type Ontology. This work is already underway with integration of cell types from the Whole Mouse Brain and Basal Ganglia taxonomies. **Figure 6D** shows an example of this integration for STRd D1 neurons^139,140^ from the primate basal ganglion and whole mouse brain taxonomies into the Cell Type Ontology. Integration with the Cell Type Ontology supports classification under a general GPi core term describing the general properties of this cell type using data from multiple references covering multiple modalities. Rationale statement records evidence for mapping the transcriptomic types in the primate and mouse atlases to the general GPi core type including annotation transfer, marker expression, and location. Each transcriptomic type includes mapping to anatomical locations based on spatial transcriptomics data and recorded using Allen Brain Cell Atlas (ABC Atlas) anatomical taxonomies, which in turn link to the general anatomy standard, Uberon, used by many platforms including CELLxGENE. Each term is linked directly to reference data on the Allen Brain Cell Atlas. Where available, CL also includes markers from BICAN taxonomies, linked to markers on the ABC atlas.

## 6. BICAN Data Release and Management

The distribution of basal ganglia cells profiled across the present studies is shown in **Figure. 7A**. Cells are anatomically presented as dorsal striatum (DS), ventral striatum (VS), and related basal ganglia areas, substantia nigra (SN), ventral tegmental area (VTA), red nucleus (RN), and subthalamic nucleus (STN). Rows are ordered by the HOMBA ontology and columns by species (top icon) and profiling technique (bottom). In addition to sn/scRNA-seq and multiome^129,141^, human cells are profiled by snm3C-seq, which jointly measure chromatin organization and DNA methylation information^142^, droplet-paired tag, which simultaneously maps histone modifications and gene expression at single-cell resolution^44^, and slide-tags, a technique in which single nuclei within an intact tissue section are tagged with spatial barcode oligonucleotides derived from DNA-barcoded beads with known position, thereby spatially localizing these profiled cells.^141^ In combination, these techniques survey the transcriptomic, epigenomic, and spatial profile of human cells. Macaque and marmoset are profiled using multiome and Patch-seq^47^ for combined profiling of electrophysiology, transcriptomics, and morphology in macaque alone. The inset shows cell totals by species and major basal ganglia divisions for a total of 17,423,201 cells, of which 92% are human, 5% macaque, and 3% marmoset.

**Figure 7.**
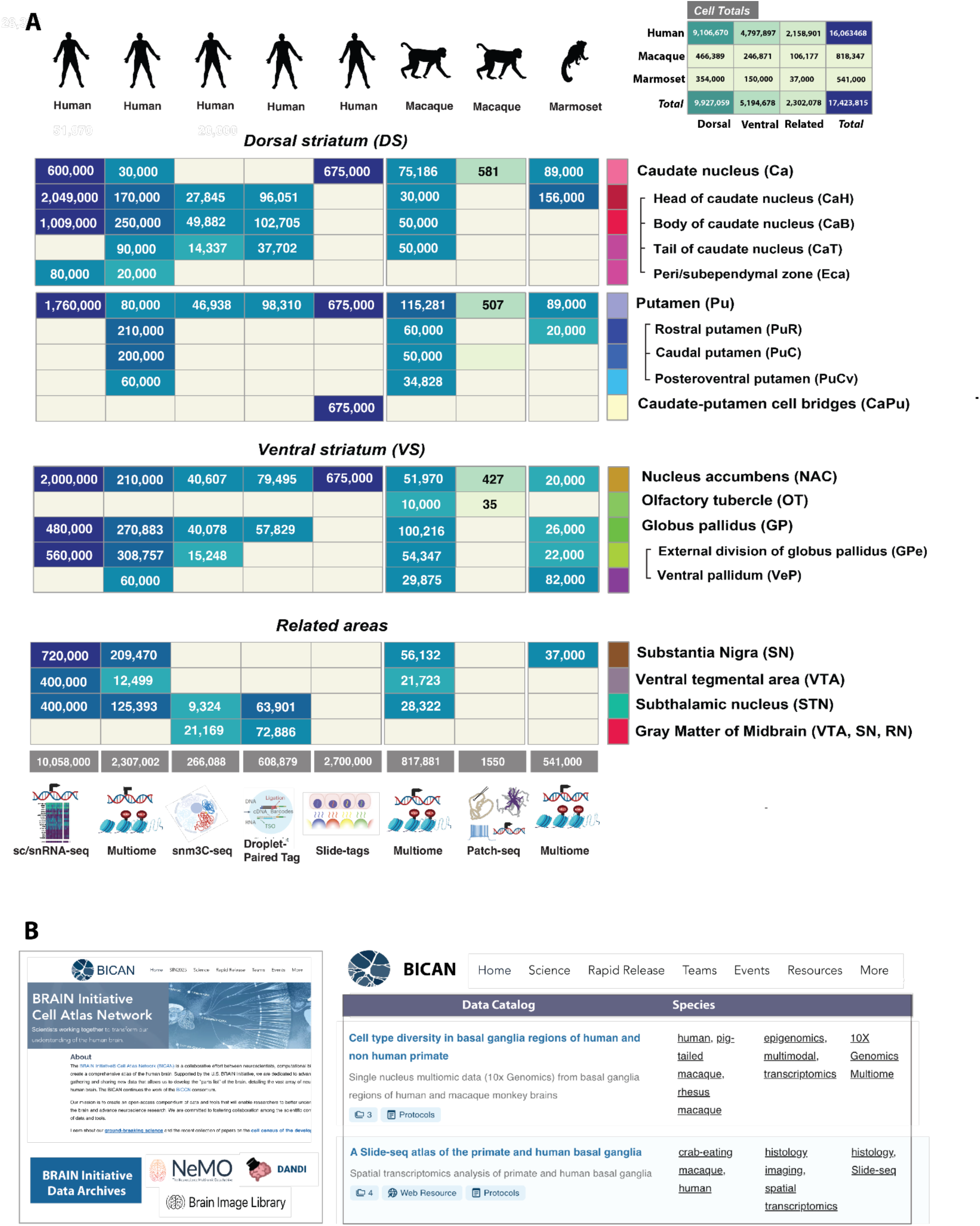
BICAN Basal Ganglia Data, Data Catalog, and Cell Inventory. A) Basal ganglia cells profiled by species, technique, and anatomic region. Rows are ordered by HOMBA ontology grouped by dorsal and ventral striatum and basal ganglia-related areas. Columns indicate species (top) and technique (bottom). Gray bar entries are cell sums for each species/technique. Upper inset shows cell totals by species and major region. B) BICAN portal home page (www.brain-bican.org) provides consortium information and access to archived data (left). Data catalog provides details of all data sets, technique summaries, web resources, and protocols.

### BICAN strategy for data release

Level 1 data includes pre-processed and processed data that have undergone at least minimal quality assurance and quality control. Pre-processed data is released by each data coordination center within 90 days^143^. For sequence-based data, datasets are disseminated after automated alignment to reference genomes and a pass or flag for release by the originating laboratory. Level 1 data are distributed in a *BICAN Rapid Release*, using federation of data and metadata across the BICAN ecosystem resources, including integration of metadata stored in NIMP, and experimental data stored at NeMO^144^, BIL^145^, and DANDI^146^ archives. The BICAN Data Catalog ^147^ (**Fig. 7B)** serves as a dashboard overview of the archives showing project-level organization across data facets such as technique, species, grant, and participating laboratory. Scientists can explore data from 6 labs, totaling 255 donors, 1461 specimens (library aliquots), and 20 data collections, accessible via a dedicated project page and specimen browser. Users may also download specimen metadata and file manifests that match the filters selected in the user interface. Analysis products tailored to individual studies will be released through the BICAN data archives. Key foundational (Level 4) data will be released for exploration and visualization via a variety of mechanisms (i.e., ABC Atlas/BKP, among others), including integration with cell type and neuroanatomical context.

### Brain Initiative Archives and BICAN

As a key part of the informatics infrastructure for the BRAIN Initiative, the data archives provide archival storage, standards, and develop software to visualize and analyze the data. The three major archives that form an essential part of the BICAN ecosystem are NeMO (RRID:SCR_016152), BIL (RRID:SCR_017272), and DANDI (RRID:SCR_017571). The archives explicitly share distribution of BICAN data where BICAN’s multimodal Patch-seq workflows: electrophysiology NWB files reside in DANDI, transcriptomic/epigenomic output is archived in NeMO, and morphological reconstructions (e.g., SWC files and image stacks) are served by BIL; cross-archive discovery occurs via the BICAN Data Catalog^148^

### Neuroscience Multi- Omic Data Archive (NeMO)

The NeMO Archive^144^ supports multiple NIH-funded consortia composed of research laboratories and sequencing centers, including the BICCN/BICAN, and Single Cell Opioid Responses in the Context of HIV Program [SCORCH; PubMed: 38879719]. The NeMO Archive has accumulated over 2.1PB of data contained in more than 8 million files, derived from sequence-based bulk and single-cell transcriptomic, methylation, and epigenomic assays, many of which are represented in **Fig.7A**. The NeMO team assists with metadata management, data ingestion and processing, and dissemination for the BICAN project. The NeMO portal allows users to explore and search for data using two methods: 1) a faceted search interface based on metadata associated with digital assets at NeMO, and 2) an advanced search feature where users can use an NCBI query-builder-like language to build a search query. The archive provides rapid release files containing Raw (Level 1) data from each BICAN laboratory as well as collections associated with publications. BICAN basal ganglia publication collections are available (NeMO ID: nemo:col-n6b0x7w0^149^) and each collection has a separate landing page with associated metadata, links to analysis files, gene counts, and relevant quality control data.

### Brain Image Library (BIL)

The BIL (RRID:SCR_017272) houses all BICAN optical microscopy data and derived secondary and tertiary data produced from optical microscopy experiments, providing web visualization and computational resources to explore the data. BIL presently contains over 5 petabytes of public data, including whole and partial brain imaging, image-based spatial transcriptomics, stained slide data, traced neurons, image-based components from multimodal experiments (such as Patch-seq), and other microscopy data modalities. The BICAN data currently in BIL consists of image-based spatial transcriptomic data (73 datasets, including MERFISH and MERSCOPE) and Patch-seq image data (273 datasets). The BIL repository provides linkage to external resources, including the BICAN NIMP resource and other BRAIN Initiative archives such as NEMO. BICAN data in BIL is tagged with the label BICAN, which can be searched for through BIL’s metadata API and web portal. Due to the inherent visual nature of data in BIL, many datasets can be directly visualized over the web. Neuroglancer and OpenSeaDragon^150^ can be used to visualize both 2- and 3-dimensional data, fluorescent microscopy data and 2D-RGB images.

### Distributed Archive for Neurophysiology Data Integration (DANDI)

DANDI is the designated archive for cellular neurophysiology and closely related imaging/behavioral data. DANDI’s use of Neurodata Without Borders (NWB)^151^, the Brain Imaging Data Structure (BIDS)^152^, and OME-Zarr ^152,153^ standards ensures interoperable description of recordings and metadata. DANDI’s holdings have expanded rapidly alongside BICAN; the archive contains ∼885 TB spread across 800+ datasets, representing ∼12,000 subjects from 21 species, with cumulative egress surpassing 8 PB in 2025. DANDI exposes data through a Web portal and a public REST API, with programmatic access via the DANDI Python/Command Line Interface for organizing, validating, uploading, and downloading; streaming methods enable analysis without full downloads; and DataLad mirrors and WebDAV support versioned, file-selective workflows. For analysis next to data, DANDI Hub (JupyterHub) provides CPU/GPU notebooks in curated environments. For interactive visualization, Neurosift and Neuroglancer are tightly integrated with DANDI’s file browser. For BICAN’s cell-type atlas efforts, DANDI is the operational home for neurophysiology and proteomic imaging.

BICAN is committed to rapid, open public release of all data it generates, and most data are freely and openly available to the public, subject only to protection of human subject identity. A subset of data has been generated from controlled access donors. Licensing is CC-BY (open use, with attribution) for non-human data, BICAN-BY-NR (open, forbids reidentification), and restricted human data accessible through the NIMH Data Archive (RRID:SCR_004434). Attribution and data citation is required for reuse, as well as compliance with laws and policies. For details, see **Suppl. Information**.

## 7. BICAN Resources and Tools

The open release of the BICAN basal ganglia data is accompanied by a rich set of tools and resources for exploring, visualizing, and analyzing the data. **Figure 8A** presents the essential toolkit that was developed through or used in the creation of these resources. The applications identified by Research Resource Identifiers (RRID) span all modalities of anatomy, single cell, and spatial data, and access BICAN atlases and CCFs (**Fig. 8B**) and ontologies (**Fig. 8C**).

**Figure 8.**
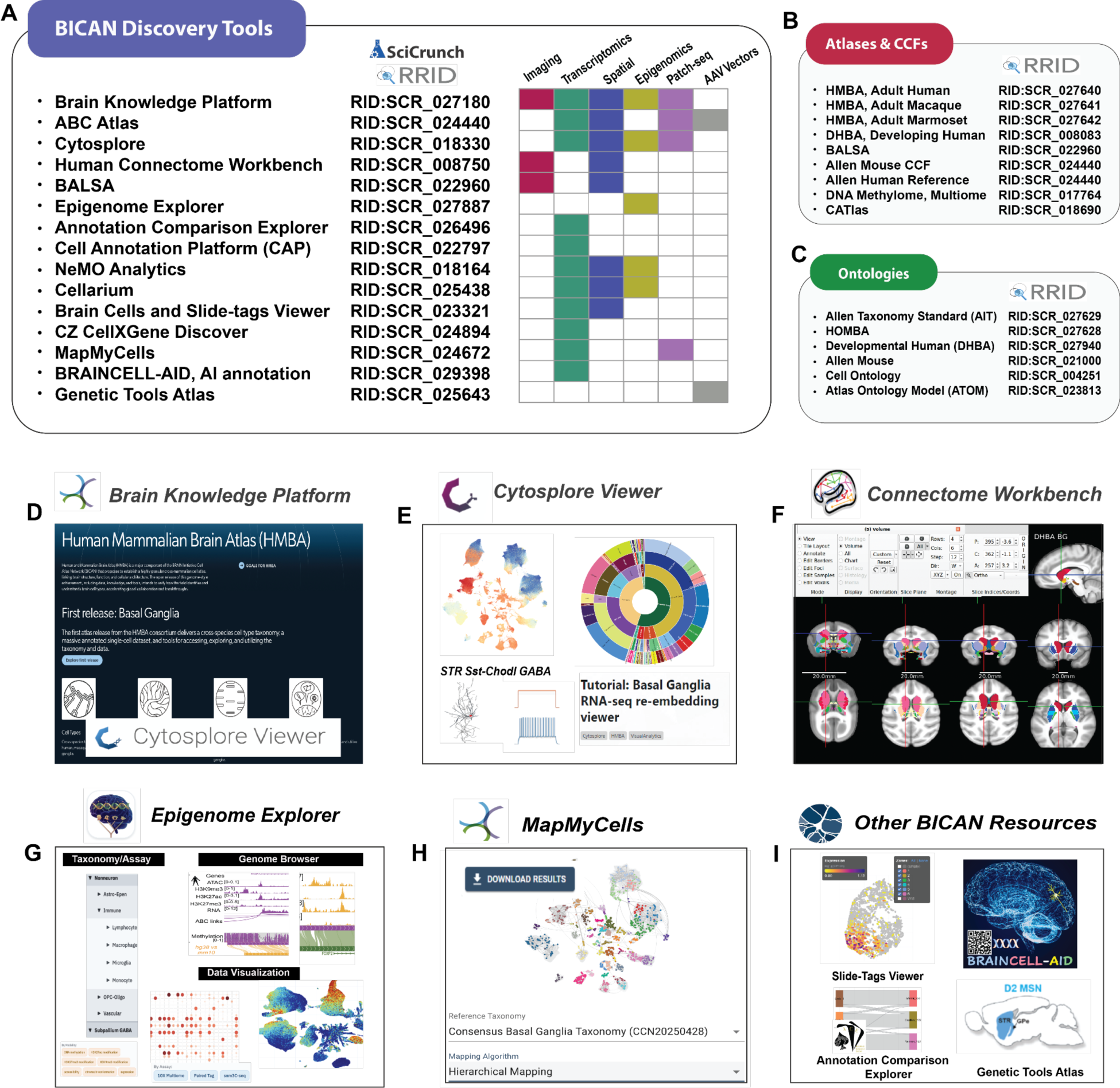
Tools and resources for BICAN basal ganglia data. A) BICAN basal ganglia-related applications, data modalities, and SciCrunch^165^ RRIDs. B, C) Basal ganglia atlases, CCFs, and ontologies. D) Brain Knowledge Platform, the primary entry point for the basal ganglia cross-species human and mammalian cell type taxonomy. E) Cytosplore, a desktop multimodal data analysis platform. F) Connectome Workbench, for exploration, anatomy and imaging data. G) The Basal Ganglia epigenome portal, for genome browser-based, multi-omics, and cross-species visualization and analyses, H) MapMyCells, for cell type mapping annotation, I) Other tools available for mapping and annotating the basal ganglia taxonomy of cell types include Slide-tags viewer^166^, BRAINCELL-AID multi-agent annotation LLM, Annotation Comparison Explorer, and Cell Annotation Platform. In collaboration with the BRAIN Armamentarium Consortium, the Genetic Tools Atlas provides cell-type-specific viral tools to target and manipulate specific cell types.

### Single Cell and Spatial Transcriptomics

The Allen Institute *Brain Knowledge Platform (BKP)* is the overarching framework for Allen Institute data, providing an open, online platform for the foundational Human and Mammalian Brain Atlas^70^ (**Fig. 8D**) including (i) the cross-species taxonomy of cell types in the human, macaque, and marmoset basal ganglia^154^, (ii) unified structural ontology, parcellations, and template images for the basal ganglia, (iii) single-cell transcriptomics, spatial transcriptomics, and Patch-seq data from the human, macaque, and marmoset basal ganglia, and (iv) tools to access, visualize, and explore the atlas, including a package to download data and metadata programmatically via Python. The repository also provides Jupyter notebooks showing examples of how to join and use these data products. As part of the BKP, the *ABC Atlas* provides viewing of gene expression and cell metadata in both single-cell and spatial transcriptomics. Four basal ganglia datasets are currently available in the ABC Atlas for exploration, each annotated with the BICAN Mammalian Basal Ganglia Consensus Cell Type Taxonomy.^155^ BICAN basal ganglia data sets are also available in CZ CELLxGENE Discover^156^ which links users to a broader corpus of data across cell atlasing communities and includes functionality for differential gene expression. *Cytosplore Viewer* and *NeMO Analytics* (see below) provide a multimodal interactive environment for data analysis and visualization. Finally, *Brain Cells and Slide-tags Data Viewer^157^* provides focused analysis of the whole mouse brain data set collected using snRNA-seq and Slide-seq spatial transcriptomics. These tools provide overlapping functionality when applied to the consensus BG taxonomy and can be selected based on the desired use case.

### Multimodal interactive data exploration

While web-based platforms provide enterprise-scale data hosting and overview visualization, they offer limited functionality for fast, interactive visual exploration that requires on-the-fly local computation across multiple datasets. *Cytosplore Viewer* (**Fig. 8E**) leverages fast local compute and compressed data access for the linked exploration of multi-modal spatial and single-cell datasets. Cytosplore’s specific focus is to complement BKP functionality with low-level detail analyses across multiple datasets, for instance: (i) Deepdives in single-cell and spatial data for in-depth re-analysis of subsets of cells, either dispatched from the ABC Atlas portal, or directly in Cytosplore Viewer. (ii) Comparative visualization for direct comparative analyses of single-cell and spatial data across multiple species and/or between brain regions. Central to these comparisons is identifying conserved and divergent feature patterns across these datasets and relating these to evolutionary and anatomical prior knowledge. (iii) Multimodal single-cell feature integration for linked visualization of, for instance, Patch-seq data (for exploration of annotated *t*-typed cells in relation to their morphological and electrophysiological features) and ATAC-seq / RNA-seq combinations. Cytosplore Viewer is a stand-alone application built on top of ManiVault Studio, a plugin-based high-performance application building environment for Windows, MacOS, and Linux.

*NeMO Analytics* (nemoanalytics.org) is a collaborative web-based system for exploring functional genomics data. The platform comprises six core components: (i) a dataset uploader/curator supporting diverse visualizations (bar, line, scatter, violin, anatomical graphics, UMAP, t-SNE, PCA); (ii) a dataset manager for organizing profiles and controlling permissions; (iii) a gene-expression browser with visualization tools, annotations, and external link-outs; (iv) interactive spatial displays; (v) a comparison tool for differential-expression analysis; and (vi) a single-cell workbench for de novo or advanced exploration of scRNA-seq data via a GUI-based, optimized *Scanpy* ^158^ pipeline. Queries by gene symbol return expression values and annotations, and cross-species comparisons of homologous genes are supported. Researchers are able to upload their own datasets and gene signatures for dissemination and exploration in this development research environment built on the gene Expression Analysis Resource (gEAR^159^).

### Accessing and Visualizing CCFs and transcriptomic samples using Connectome Workbench and BALSA

The Connectome Workbench platform^160^ (**Fig. 8F**) enables visualization and analysis of human, macaque, and marmoset brain templates, atlases, and CCFs, including surface-based representations of cerebral cortex that respect the topology of the cortical sheet and volume-based representations of subcortical structures. To enable rapid and flexible navigation between putative homologous parcels (or their parent structures) across species, Connectome Workbench indexes annotated areas of nomenclature-harmonized parcellations (e.g., SARM, D99, SAM, DHBAv2) into the HOMBA ontology. The ontology preserves the provenance of each area and can accommodate additional areal metadata in the flexible JSON format. Parcellations from previously published SARM, D99, and SAM atlases were transformed into the same coordinate spaces as their species-specific counterparts currently being annotated with HOMBA on native templates. Our templates consist of the existing HCP3T1071 in MNI space for human, plus novel state-of-the-art non-human primate MRI templates with HCP-style image acquisition and preprocessing: Mac25Rhesus_v5, Mac25Cyno_v3, and MarmosetRIKEN25_v1 for rhesus macaque, cynomolgus macaque, and common marmoset, respectively. MRI-based templates and parcellations are accessible in the Brain Analysis Library of Spatial Maps and Atlases (BALSA) neuroimaging study results repository^161^, with the individual-level HCP data from which they were derived accessible in ConnectomeDB powered by BALSA. The initial CCF release emphasizes the basal ganglia and associated subcortical structures and includes volumetric representations; however, surface-based representations will be incorporated for cortical structures in the future.

### Exploration of Epigenetics

The *Basal Ganglia Epigenome Portal ^162^* (RRID:SCR_027887; **Fig. 8G**) contributes an integrative resource for exploring epigenetic regulation in the human brain by providing a web-based platform that unifies single-cell transcriptomic and epigenomic data from the basal ganglia. By combining chromatin accessibility, DNA methylation, histone modifications, chromatin conformation, and gene expression^71,162^ within a genome browser–centered framework, the resource enables intuitive, cell-type-resolved interrogation of regulatory mechanisms. Its support for cross-species comparative analysis and interactive visualization exemplifies how epigenomic data integration can facilitate discovery and broaden access to complex multiomic datasets within the research community.

### Mapping and Annotating Cell Types

A variety of tools are available for mapping and annotating the basal ganglia taxonomy of cell types (**Fig. 8**). Single-cell mapping methods transform raw, heterogeneous single-cell datasets into interpretable, comparable, and biologically grounded information. Among many methods for aligning single cell data to reference taxonomies, *MapMyCells* (**Fig. 8H**)^163^ provides out-of-the-box capability to map onto the growing set of high-quality cell type taxonomies including the HMBA cross-species consensus basal ganglia. As an open-source library, MapMyCells provides tools to map to any hierarchical cell type taxonomy given minimal preprocessing with functions provided by MapMyCells itself. MapMyCells is deterministic, scalable, and provides a readily interpretable metric of confidence in the resultant mappings. *Annotation Comparison Explorer^164^* provides an interactive tool for aligning cell type nomenclature and associated metadata across different cell type taxonomies created by the Allen Institute, BICAN, and Alzheimer’s disease research communities. The *Cell Annotation Platform^126^*plays a critical role in visualizing gene expression data and cell metadata for the primary BG taxonomy itself, and more importantly acts as the central database for tracking all community cell type annotations. Essentially, a community annotator can use CAP to visualize gene expression data, select a cell type to provide feedback on, select a type of annotation (e.g., agree, split, merge, other), and then provide comments and associated context therein. All feedback from all users is stored in a central database which can be exported by cell type taxonomy owners for integration in future taxonomy versions. As an additional resource for integrated cell type annotation, *Cellarium (RRID:SCR_025438) Cell Annotation Service (CAS)*, co-developed by 10X Genomics and Broad Institute Data Sciences Platform^167^, is available in the form of Python Client Library and API. CAS builds a vector search index derived from low-dimensional embeddings of a comprehensive repository of publicly available single-cell transcriptomics data including CZI CellXGene data catalog and uses machine learning algorithms for more seamless exploration of vast single-cell datasets. It provides a cloud search engine where users can submit queries to receive rapid and accurate annotation for single-cell omics data.

### Large Language Models and Cell Types

Single-cell RNA-Sequencing technologies have revolutionized brain cell type identification, yet functional annotation of novel or rare cell types remains difficult due to incomplete markers and fragmented literature. While large language models (LLMs) show promise, their accuracy is limited by factual inconsistencies. BICAN developed Brain Cell type Annotation and Integration using Distributed AI (BRAINCELL-AID) ^13,168^, a multi-agent AI system combining LLM fine-tuning, literature mining, and retrieval-augmented generation (RAG)^169^ for accurately annotating brain marker gene sets and cell types. Trained on 7,000+ gene sets from MSigDB^170^, BRAINCELL-AID achieved accuracies of 77% and 74% on mouse and human datasets, respectively, outperforming state-of-the-art methods. BRAINCELL-AID generates biologically grounded annotations for brain cells and offers a community resource for expert evaluation and collaborative annotation for BICAN data.

### Genetic Tools Atlas

The BRAIN Armamentarium Consortium has developed an extensive collection of cell-type-specific viral tools designed to target and manipulate specific cell types. These tools were validated with mapping to the HMBA consensus BG taxonomy and implemented for discovery work in BICAN projects.^122^ Using bulk or single-nucleus ATAC-seq and multiomic data, cell-type-specific gene regulatory elements are identified, particularly enhancers, and linked to the cell types identified through snRNA-seq. These efforts culminate in the *Cross-species Enhancer Ranking Pipeline (CERP),* a robust computational pipeline for identifying successful enhancers with a high degree of accuracy. This pipeline identifies and prioritizes testing of promising candidate cell type enhancers for targeting homologous cell types across mammalian species. This viral tool collection comprises hundreds of enhancer adeno-associated virus (AAV) vectors, including more than 400 basal ganglia cell type enhancers, each with high-resolution mouse brain expression images in the *Genetic Tools Atlas*. ^171^ The most promising tools were also subjected to detailed molecular validation. These collective resources provide researchers with a powerful toolkit for targeting and studying the diverse cell types of the basal ganglia across multiple species to advance our understanding of BG function and dysfunction in health and disease.

## 8. Toward a Foundational Cell Type Atlas of the Human Brain

The BICAN consortium has taken a first major step toward a foundational whole-brain human cell atlas, establishing a framework for defining and organizing brain cell types and their states across the brain and in relation to model organisms. Foundational efforts already outline coarse cell-type organization across the human brain^31,55^ and provide high-resolution molecular characterization of the adult mouse brain^13^, demonstrating the feasibility of comprehensive cellular mapping. Centered on the basal ganglia, the present multi-species, multimodal atlas provides a scalable framework for extending these approaches to the entire human brain. Completion of such a reference will unify how the field defines, compares, and interprets brain cell types, analogous in role to the human genome as a shared, high-quality reference for the community.

The long-term objective of BICAN is to construct a unified reference that (i) maps genomically defined cell types across the entire human and non-human primate brain, (ii) aligns homologous cell types across species, (iii) links cellular identity to structural and functional architecture, (iv) resolves spatial distributions of cell types, (v) integrates morphoelectric and physiological properties, and (vi) quantifies robustness and variation across individuals. The present basal ganglia atlas represents the first realization of these goals, delivering high-resolution, cross-species characterization across human, macaque, and marmoset.

At the core of this effort is the HMBA, which integrates a cross-species cell-type taxonomy with harmonized anatomical reference frameworks. Anchored in single-nucleus transcriptomics and complemented by epigenomic and spatial profiling, HMBA establishes data-driven linkages between molecular identity and anatomical context. This work is synergistic with broader initiatives such as the Human Cell Atlas^31^, collectively advancing comprehensive, interoperable references of human cellular organization across tissues and states.

Integration of diverse single-cell and spatial genomics datasets within this framework has yielded new insights into basal ganglia structure and function. These include the construction of a unified multispecies taxonomy^24^, identification of conserved and species-specific cell types, and discovery of previously unrecognized mesoscale organization, including spatial gradients^84^, striosome–matrix compartmentalization^85^, specialized cellular communities that link molecular identity with anatomical and circuit context^86^, and molecular zonation of the human striatum^85^. Analyses of inter-individual variation revealed widespread, cell-type-specific transcriptional changes associated with aging, with comparatively limited sex-associated effects, and remarkably demonstrates that age can be predicted from gene expression profiles across multiple cell types^87^. Complementary multimodal studies extend these findings into the regulatory domain, integrating chromatin accessibility, histone modification, DNA methylation, three-dimensional genome organization^23^, and gene expression to define the regulatory architecture of brain cell types.

Despite this progress, significant challenges remain in establishing a comprehensive whole-brain reference. A robust donor and sampling strategy is essential, and these efforts now need to be extended to the entire brain. BICAN employs systematic spatial sampling across brain regions with sufficient replication to quantify inter-individual variability, guided by the HOMBA ontology to ensure consistent anatomical annotation. Advances in standardized neuropathological assessment and metadata capture used by the consortium and captured by the NIMP sampling and sequencing portal have been essential to contextualize molecular data and support downstream interpretation. Without this rigor, apparent cell types risk reflecting sampling artifacts rather than biological reality. Equally important is the harmonization of tissue processing and assay pipelines. Standard operating procedures for tissue handling, cell and nuclei isolation, library preparation, sequencing, and data processing, implemented across shared platforms and coordinated centers, ensure consistency and comparability of datasets. These standardized workflows enable mapping of data generated across sites into a common reference framework, supported by shared data models and interoperable infrastructure.

Recent advances in artificial intelligence, including deep learning and large language models, offer new opportunities to enhance the analysis and interpretation of large-scale biological data. A BICAN AI Interest Group promotes integration of these methods to improve data FAIR-ness and analytical capability. AI techniques are increasingly playing an important role in the interpretation and analysis of single-cell data. However, it is important to address ethical issues arising from generative AI and LLMs—including lack of transparency and explainability in models, biases in training data and resultant models, and the inaccessibility of computational resources for employing AI techniques across institutions.

Realization of a whole-brain atlas as a durable community resource requires sustained governance and release engineering. The atlas must be versioned, with transparent provenance, reproducible pipelines, and clear documentation of updates. A coordinated infrastructure supports data management, integration, and dissemination, ensuring alignment between investigators and public archives. Importantly, the atlas should be viewed not as a static product but as a living reference system—continuously refined through community contribution, extension to new species and modalities, and incorporation of emerging data. Ethical considerations, including consent, privacy, and controlled data access, must be embedded within this framework. Together, these efforts establish a FAIR, extensible ecosystem that enables researchers to map, compare, and interpret data within a shared cellular reference framework, providing the foundation for scaling from regional atlases to a comprehensive, whole-brain representation of human cell-type organization and function.

## Supporting information

BICAN Resources

## Acknowledgements

This publication was supported by and coordinated through the Brain Initiative Cell Atlas Network (BICAN). Research reported in this publication was supported by the National Institutes of Health (NIH) BRAIN Initiative under award numbers UM1MH134812, UM1MH130994, UM1MH130981, UM1MH130966, UF1MH128339, U24NS133077, U24MH130988, U24MH130968, U24MH130919, U24MH130918, U01MH130995, U01MH130962, RF1MH133777, R24MH117295, R24MH114793, R24MH114788, F32MH138113, UM1MH130991, U01MH130907, R01MH138332, R01MH134833, R01MH134809, R01MH134800, R01MH134799, R24MH114785, U01MH124619, U01MH124602, U01MH117079, U01MH117072, U01MH117023, U01MH116990, U01MH114828, RF1MH128970, RF1MH128969, RF1MH128949, RF1MH128888, RF1MH128885, RF1MH128876, RF1MH128875, RF1MH128867, RF1MH128866, RF1MH128861, RF1MH128842, RF1MH128841, RF1MH128840, RF1MH128838, RF1MH124612, RF1MH124606, RF1MH124605, RF1MH124598. Research in this publication was also supported by the Netherlands Organization for Scientific Research NWO:024.004.012, Japan Society for the Promotion of Science (JSPS) JP22H0492, and Japan Agency for Medical Research and Development (AMED) JP23wm0625001 and JP18dm0307006

## Authors and Contributions

### BICAN Contributing Principal Investigators

M. Margarita Behrens,Timothy E. Brown, Joseph R. Ecker, Bruce Fischl, Winrich A. Freiwald, Satrajit S. Ghosh, Thomas J. Grabowski, Michael Hawrylycz, Takuya Hayashi, Rebecca Hodge, Bing-Xing Huo, C. Dirk Keene, Henry Kennedy, Christiaan P.J. de Kock, Arnold Kriegstein, Fenna M. Krienen, Boudewijn Lelieveldt, Ed Lein, Daofeng Li, Sten Linnarsson, Chongyuan Luo, Evan Macosko, Huibert D. Mansvelder, Maryann E. Martone, Steve McCarroll, Shoaib Mufti, Lydia Ng, David Osumi-Sutherland, Mercedes Paredes, Carol Thompson, Timothy Tickle, Andreas S. Tolias, Bing Ren, Alexander J. Ropelewski, Gabor Tamas, David C. Van Essen, Alan M. Watson, Owen White, Hua Xu, Xiangmin Xu, Hongkui Zeng, W. Jim Zheng, Guo-Qiang Zhang

### Manuscript Writing, Figure Generation

Ashwin Bhandiwad, Timothy E. Brown, Song-Lin Ding, Mathew F. Glasser, Satrajit S. Ghosh, Michael Hawrylycz, Rebecca Hodge, Hao Huang, Bing-Xing Huo, Juan Eugenio Iglesias, Nelson J. Johansen, Christiaan P.J. de Kock, Ed Lein, Boudewijn Lelieveldt, Daofeng Li, Huibert D. Mansvelder, Maryann E. Martone, Jeremy A. Miller, David Osumi-Sutherland, Patrick Ray, Andrea D. Rivera, Burke Q. Rosen, Timothy Tickle, Carol Thompson, David C. Van Essen, Owen White, W. Jim Zeng, Guo-Qiang Zhang

### Manuscript Editing

Trygve E. Bakken, M. Margarita Behrens, Wubin Ding, Satrajit S. Ghosh, Michael Hawrylycz, Yasmeen Hussain, Juan Eugenio Iglesias, Nelson J. Johansen, Daofeng Li, Maryann E. Martone, Partha P. Mitra, Mercedes Parcedes, Burke Q. Rosen, Carol Thompson, Jonathan T. Ting, David C. Van Essen, Ting Wang, Morgan Wirthlin, Owen White, Hongkui Zeng, Guo-Qiang Zhang, Hongkui Zeng, Wenjin Zhang

### Human and Mammalian Brain Atlas

Trygve E. Bakken, Ashwin Bhandiwad, Jennie Close, Tanya L. Daigle, Rachel Dalley, Song-Lin Ding, Yuanyuan Fu, Aaron Garcia, Tim Jarsky, Jimena Garcia, Mathew F. Glasser, Michael Hawrylycz, Takuya Hayasahi, Madeleine N. Hewitt, Rebecca Hodge, Yasmeen Hussain, Juan Eugenio Iglesi, Takuro Ikenda, Tomoko Ishibuchi, Fenna M. Krienen, Nelson J. Johansen, Brian E. Kalmbach, Henry Kennedy, Christiaan P.J. de Kock, Brian R. Lee, Ed Lein, Sten Linnarsson, Xiao-Ping Liu, Brian Long, Huibert D. Mansvelder, Delissa McMillan, Jeremy A. Miller, Lydia Ng, Philip Mardoum, Tyler Mollenkopf, Shoaib Mufti, Takayuki Ose, Michael Platt, David Osumi-Sutherland, Raymond Sanchez, Burke Q. Rosen, Mathew Schmitz, Stephanie Seeman, Kimberly A. Smith, Noah Snyder-Mackler, Staci Sorensen, Gabor Tamas, Jonathan T. Ting, Andreas S. Tolias, Meghan Turner, David C. Van Essen, Morgan Wirthlin, Zizhen Yao, Faraz Yazdani, Dan Yuan, Hongkui Zeng

### Brain Specimens Core

Virginia Allhusen, Derick Aranda, Sabina Berretta, Jillian V. Berry, Thomas Blanchard, H Mark Brooks, William E. Bunney, Preston Cartagena, Daniel Child, John Crawford, Winrich Freiwald, Thomas J. Grabowski, Mathew G. Heffel, Vahram Harry Haroutunian, Elizabeth Head, C. Dirk Keene, Jason M Knight, Caitlin Latimer, David A. Lewis, Christine MacDonald, Firoza Mamdani, Edwin Monuki, Stefano Marencos,Rashed Nagra, Amber Nolan, Hannah Lui Park, William K. Scott, Pedro Adolfo Sequeira, Craig Stark, Sabrina Stein, Xiaoyan Sun, Bryan M. Tran, Lorena Valenzuela, Xiangmin Xu, Faraz, Yazdani, William Yong

### Atlas of Human Brain Variation

Kiku Ichihara, Evan Macosko, Steve McCarroll, Charles Vanderburg

### Center for Multiomic Human Brain Cell Atlas

Cesar Barragan, Anna Bartlett, Cindy T. Baez Becerra, M. Margarita Behrens, Eric Boone, Rosa G Castanon, Lei Chang, Wubin Ding, Joseph R. Ecker, Jesus Flores, Amit Klein, Kai Li, Colin Kern, Daofeng Li, Alexander Monell, Joseph R. Nery, Jacquelin Olness, Emma Osgood, William Owens, Bing Ren, Jonathan Rink, Chu-Yi Tai, Ting Wang, Yang Xie, Xiangmin Xu, Wenjin Zhang, Zoey Zhao, Guojie Zhong, Quan Zhu

### Epigenetic and Chromosomal Architecture of Developing Human Brain

Mathew G. Heffel, Chongyuan Luo, Oier Pastor-Alonso, Mercedes F. Paredes

### Multidisciplinary Center for Developing Human and NHP Brain Cell Atlases

Hao Huang. Arnold Kriegstein

### NIMP, NIMP Analytics and APIs

Rashmie Abeysinghe, Antarr Byrd, Wei-chun Chou, Licong Cu, Erge Edgu-Fry, Yan Huang, Xiaojin Li, Shiwei Lin, Kimberly A. Smith, Shiqiang Tao, Ling Tong, Guo-Qiang Zhang

### Imaging and CCF

Joonas Autio, Ashwind Bhandiwad, Timothy S. Coalson, Song-Lin Ding, Bruce Fischl, Aaron Garcia, Mathew F. Glasser, Thomas J. Grabowski, Takuya Hayashi, Hao Huang, Mike Huang, Yujie Hou,Takuro Ikeda, Juan Eugenio Iglesias, Tomoko Ishibuchi, Camille Lamy, Henry Kennedy, Lydia Ng, Pierre Misery, Masahiro Ohno, Takayuki Ose, Burke Q. Rosen, Akiko Uematsu, David C. Van Essen, Julien Vezoli

### NeMO archive

Seth Ament, Robert Carter, Victor Felix, Chakshu Gandhi, Michelle Giglio, Brian Herb, Theresa Hodges, Olukemi Ifeonu, Roland Laboulaye, Jonathan Lim, Anup Mahurkar, Suvarna Nadendla, Dustin Olley, Joseph Receveur, William Scerbo, Mike Schor, Akira Watanabe, Owen White

### BIL archive

Mariah Kenney, Alexander J. Ropelewski, Iaroslavna Vasylieva, Alan M. Watson,

### DANDI archive

Cody Baker, Roni Choudhury, Nima Dehghani, Benjamin Dichter, Satrajit S. Ghosh, Yaroslav O. Halchenko, Kabilar Gunalan, Dorota Jarecka, Austin Macdonald, Jeremy Magland, Jacob Nesbitt, Isaac To, Michael VanDenburgh, John T. Wodder II

### Brain Cell Data Center

Katherine Baker, Pamela M. Baker, Anita Bandrowski, Edyta Vieth, Prajal Bishwakarma, Elysha Fiabane, Tim Fliss, Limary Gonzalez, Michael Hawrylycz, Shanshan Christine Lin, Maryann E. Martone, Sven Otto, Patrick Ray, Carol Thompson

### Ontology and Data Standards

Rashmie Abeysinghe, Pamela M. Baker, Anita Bandrowski, Licong Cui, Song-Lin Ding, Na Hong, Yan Huang, Xiaojin Li, Maryann E. Martone, Jeremy A. Miller, David Osumi-Sutherland, Patrick Ray, Andrea D. Rivera, Kimberly A. Smith, Shiqiang Tao, Carol Thompson, W. Jim Zheng, Hua Xu, Guo-Qiang Zhang

### Data Infrastructure

Jason Alexander, Cameron Bielstein, Kaitlyn Colbert, Peter DiValentin, Tim Dolbeare, Yan Huang, Zachary Madigan, Imran Majeed, Jessica Malloy, Phillip Mardoum, Christopher B. Morrison, Shoaib Mufti, Cade Reynoldson, Raymond Sanchez, Lane Sawyer, Shiqiang Tao, Carol Thompson, Timothy Tickle, Shane Vance, Guo-Qiang Zhang

### Data Science Platform

Aseel Awdeh, Mehrtash Babadi, Robert Sidney Cox III, Stephen Fleming, Fedor Grab, Giles Hall, Bing-Xing Huo, Farzaneh Khajouei, Elizabeth Kiernan, Anton Kovalsky, Nikelle Mirabito, Timothy Tickle, Erin Schoenbeck, Jessica Way, Yang Xu

### Software Tools and Resources

Tek Raj Chhetri, Timothy S. Coalson, Scott F. Daniel, Matt F. Glasser, Na Hong, Bing-Xing Huo, Daofeng Li, Boudewijn Lelieveldt, Jeremy A. Miller, Tyler Mollenkopf, Shoaib Mufti, Burke Q. Rosen, David Osumi-Sutherland, Puja Trivedi, Ting Wang, Yimin Wang, Hua Xu, W. Jim Zheng, Wenjin Zhang,

### Neuroethics

Timothy E. Brown, Zoë J. Hale, Maria Sourdi

### Project Management

Yasmeen Hussain, Lauren Kruse, Sam Caldwell, Xiaojin Li, Susan Sunkin, Shiqiang Tao, Carol Thompson, Guo-Qiang Zhang

