## Supplementary material for "A Community Standard Multispecies Cell Atlas of the Basal Ganglia": BICAN Resources

### BICAN Resource Inventory

| Resource | RRID / PID | Description | Usage Notes |
|---|---|---|---|
| **Analysis Pipelines** | | | |
| **ATAC Pipeline** | RRID:SCR_024656 | An open-source, cloud-optimized pipeline developed in collaboration with members of the BRAIN Initiative (BICCN and BICAN Sequencing Working Group) and SCORCH. It supports the processing of 10x single-nucleus data generated with 10x Multiome ATAC-seq (Assay for Transposase-Accessible Chromatin), a technique used in molecular biology to assess genome-wide chromatin accessibility. | |
| **BuildIndices Pipeline** | RRID:SCR_027630 | An open-source, cloud-optimized pipeline developed in collaboration with BICCN and BICAN. The workflow filters GTF files for selected gene biotypes, calculates chromosome sizes, and builds reference bundles with required files for STAR and bwa-mem2 aligners. | |
| **Multiome Single-cell ATAC and Gene Expression Pipeline** | RRID:SCR_024217 | An open-source, cloud-optimized pipeline developed in collaboration with members of the BRAIN Initiative (BICCN and BICAN Sequencing Working Group) and SCORCH. It supports the processing of 10x 3' single-cell and single-nucleus gene expression (GEX) and chromatin accessibility (ATAC) data generated with the 10x Genomics Multiome assay. | |
| **Optimus Pipeline** | RRID:SCR_018908 | An open-source, cloud-optimized pipeline developed by the Data Coordination Platform (DCP) of the Human Cell Atlas (HCA) Project as well as BICCN. It supports the processing of any 3' single-cell and single-nucleus count data generated with the 10x Genomics v2 or v3 assay. | |
| **Paired-Tag Pipeline** | RRID:SCR_025042 | An open-source, cloud-optimized pipeline developed in collaboration with BICCN and BICAN. It supports the processing of 3' single-nucleus histone modification data (generated with the paired-tag protocol) and 10x gene expression (GEX) data generated with the 10x Chromium Multiome assay. | |
| **Single Nucleus Methyl-Seq and Chromatin Capture (snm3C) Pipeline** | RRID:SCR_025041 | An open-source, cloud-optimized computational workflow for processing single-nucleus methylome and chromatin contact (snm3C) sequencing data. The workflow is designed to demultiplex and align raw sequencing reads, call chromatin contacts, and generate summary metrics. | |
| **Slide-Seq Pipeline** | RRID:SCR_023379 | An open-source, cloud-optimized pipeline developed in collaboration with BICCN and BICAN. It supports the processing of spatial transcriptomic data generated with the Slide-seq (commercialized as Curio Seeker) assay. | |
| **Slide-tags Pipeline** | RRID:SCR_027567 | An open-source, cloud-optimized workflow for processing spatial transcriptomics data. It supports data derived from spatially barcoded | |

| Resource | RRID / PID | Description | Usage Notes |
|---|---|---|---|
| | | sequencing technologies, including Slide-tags-based single-molecule profiling. The pipeline processes raw sequencing data into spatially resolved gene expression matrices, ensuring accurate alignment, spatial positioning, and quantification. | |
| **Smart-seq2 Single Nucleus Multi-Sample Pipeline** | RRID:SCR_021312 | Pipeline developed in collaboration with BICCN to process single-nucleus RNAseq (snRNAseq) data generated by Smart-seq2 assays. The pipeline's workflow is written in WDL, is freely available in the WARP repository on GitHub, and can be run by any compliant WDL runner (e.g. Cromwell). | |
| **Atlases & Coordinate Frameworks** | | | |
| **Allen Adult Mouse CCF, Stereotaxic (Wang 2020)** | RRID:SCR_020999 | Mouse Common Coordinate Framework at 10µm voxel resolution created from a group average of 1,675 young adult C57Bl6/J mice imaged by serial two-photon tomography, with the entire brain parcellated directly in 3D, labeling every voxel with a brain structure spanning 43 isocortical areas and their layers, 314 subcortical gray matter structures, 81 fiber tracts, and 8 ventricular structures. | |
| **Allen Brain Cell Atlas (ABC)** | RRID:SCR_024440 | A high-resolution transcriptomic and spatial cell-type atlas across the entire mouse brain, integrating several whole-brain single-cell RNA-sequencing (scRNA-seq) datasets containing a total of ~4 million cells passing rigorous quality-control (QC) criteria. | *Basal ganglia data in the ABC atlas* |
| **Allen Human Reference Atlas, 3D (2020)** | RRID:SCR_017764 | A parcellation of the adult human brain in 3D, labeling every voxel with a brain structure spanning 141 structures, drawn on the MRI reference brain volume "ICBM 2009b Nonlinear Symmetric," a non-linear average of the MNI152 database of 152 normal brain images. | |
| **Allen Institute Atlas Primer** | — | Provides a single point of access to all HMBA CCFs, including links to Neuroglancer visualizations. | |
| **Brain Analysis Library of Spatial maps and Atlases (BALSA)** | RRID:SCR_022960 | A database for hosting and sharing neuroimaging and neuroanatomical datasets for human and primate species. | Templates and HMBA BG annotations for human, cynomolgus macaque, rhesus macaque, and common marmoset as well as a HOMBA /json are available on BALSA at: https://balsa.wustl.edu:443/study/view/ggDBg?token=Mj6Dm |
| **Developing Human Brain Atlas (DHBA)** | RRID:SCR_008083 | Atlas of developing human brain for studying transcriptional mechanisms involved in human brain development. Consists of RNA sequencing and exon microarray data profiling up to sixteen cortical and subcortical structures across the full course of human brain development, high resolution neuroanatomical transcriptional profiles, in situ hybridization image data, and reference atlases with supporting histology, MRI and DWI data. | *Ding, S.-L. et al. Comprehensive cellular-resolution atlas of the adult human brain. J Comp Neurol 524, 3127–3481 (2016)* |

| Resource | RRID / PID | Description | Usage Notes |
|---|---|---|---|
| **HMBA Adult Human Brain Atlas** | RRID:SCR_027640 | Human Common Coordinate Framework based upon HCP young adult group averaged MRI template in MNI 6th generation asymmetric nonlinear space. The initial version is annotated with the basal ganglia structures from the Harmonized Ontology of the Mammalian Brain Anatomy. | *Basal ganglia in HMBA Adult Human Brain Atlas* |
| **HMBA Adult Macaque Brain Atlas** | RRID:SCR_027641 | Macaque Common Coordinate Framework based upon Mac25Rhesus group averaged MRI template, annotated with structures aligned with the Harmonized Ontology of the Mammalian Brain Anatomy. | |
| **HMBA Adult Marmoset Brain Atlas** | RRID:SCR_027642 | Marmoset Common Coordinate Framework based upon Marmoset Riken25v1 adult group averaged MRI template, annotated with structures aligned with the Harmonized Ontology of the Mammalian Brain Anatomy. | |
| **Human Connectome Young Adult Template** | — | High-resolution 3T MR scans from young healthy adult twins and non-twin siblings (ages 22–35) using four imaging modalities: structural images (T1w and T2w), resting state fMRI, task fMRI, and diffusion imaging. | |
| **BICAN Infrastructure** | | | |
| **BICAN Data Catalog** | RRID:SCR_027884 | Catalog of BICAN datasets with links to primary data within the archives. | *Basal ganglia data package* |
| **BICAN Portal** | — | Main website for the BICAN project. | |
| **Brain Knowledge Platform** | RRID:SCR_027180 | Comprehensive multi-species platform for neuroscience teams to understand brain cells and anatomy. Includes catalog of relevant data and projects, as well as visualization tools for understanding brain cell types. | |
| **Broad Institute Genomics Platform** | RRID:SCR_027987 | Large-scale research facility that generates, analyzes, and interprets high-throughput genomic data to understand the genetic basis of disease. | *BICAN sequencing center* |
| **Broad Terra** | RRID:SCR_021648 | An open-source, cloud-native platform developed by the Broad Institute in collaboration with Verily and Microsoft, designed for biomedical researchers to access, analyze, and share large-scale data securely. | *Cloud-optimized pipelines used in BICAN for processing biological data* |
| **New York Genome Center** | — | An independent, non-profit academic research institution based in New York City, founded in 2011. Acts as a collaborative hub for genomics research, focusing on advancing clinical diagnostics, genomic technologies, and data analysis. | *BICAN sequencing center* |
| **NHash Identifier** | RRID:SCR_025313 | Lightweight, blockchain-style resource identifiers for tracking research resource linkage, provenance, utilization, and visualization, generated automatically in a completely distributed fashion with virtually no risk for identifier collision. | |

| Resource | RRID / PID | Description | Usage Notes |
|---|---|---|---|
| **NIH NeuroBioBank** | RRID:SCR_003131 | A national resource for investigators utilizing human post-mortem brain tissue and related biospecimens for their research to understand conditions of the nervous system. | |
| **NIMP: Neuroanatomy-anchored Information Management Platform** | RRID:SCR_024684 | Centralized system for tracking specimen and sequencing workflows in BICAN. NIMP consists of two portals: the Specimen Portal for tissue management from donors to brain slabs and annotated brain samples, and the Sequence Library (SeqLib) Portal for managing the workflow from tissue to data deposition to assay-dependent, data-modality-specific archives. | |
| **Data Archives & Repositories** | | | |
| **Brain Image Library (BIL)** | RRID:SCR_017272 | A platform for publishing, sharing, and visualizing large light microscopic 2- and 3-D brain datasets. | *Access BICAN data in BIL via the BICAN Data Catalog* |
| **Distributed Archives for Neurophysiology Data Integration (DANDI)** | RRID:SCR_017571 | A platform for publishing, sharing, and processing neurophysiology data funded by the BRAIN Initiative. | *Access BICAN data in DANDI via the BICAN Data Catalog* |
| **National Institute of Mental Health Data Archive (NDA)** | RRID:SCR_004434 | Data archive hosted by NIMH making available human subjects data collected from hundreds of research projects across many scientific domains. NDA provides infrastructure for sharing research data, tools, methods, and analyses enabling collaborative science and discovery. De-identified human subjects data, harmonized to a common standard, are available to qualified researchers. | *Houses de-identified BICAN data not consented for open sharing* |
| **Neuroscience Multi-Omic Archive (NeMO)** | RRID:SCR_016152 | Data repository specifically focused on storage and dissemination of omic data generated from BRAIN Initiative and related brain research projects. | *Basal ganglia dataset collection in NeMO* |
| **Reference Taxonomies** | | | |
| **HMBA Cross-species Consensus Taxonomy of Basal Ganglia** | — | The consensus basal ganglia cell type taxonomy resulting from iterative clustering and cross-species integration of transcriptomic data from single-nucleus 10x Genomics multiomic profiling. The taxonomy encompasses neurons from key structures within the basal ganglia, including the caudate (Ca), putamen (Pu), nucleus accumbens (NAc), the external and internal segments of the globus pallidus (GPe, GPi), ventral pallidus (VeP), subthalamic nucleus (STN), and substantia nigra (SN). | *Johansen et al. (2025)* |
| **Ontologies & Taxonomies** | | | |
| **Allen Institute Taxonomy Standard (AIT)** | RRID:SCR_027629 | Aligned taxonomy format. | |

| Resource | RRID / PID | Description | Usage Notes |
|---|---|---|---|
| **Atlas Ontology Model** | RRID:SCR_023813 | A standardized, machine-readable framework designed to harmonize brain atlases by defining the relationships between four core elements: reference data, coordinate system, annotation set, and terminology. | *Kleven, H. et al. Sci Data 10, 486 (2023)* |
| **BICAN Basal Ganglia Nomenclature Standard** | — | Cell type naming standard for basal ganglia transcriptomic cell types. | *Johansen et al. (2025)* |
| **Cell Ontology** | RRID:SCR_004251 | A reference ontology designed to classify and describe cell types across different organisms. It serves as a resource for model organism and bioinformatics databases. | *Basal Ganglia Cell Ontology in Cell Ontology* |
| **Developmental Human Brain Atlas Ontology (DHBA)** | RRID:SCR_027940 | An application ontology built by combining ontologised versions of the Allen Institute Developing Human Brain Atlas (DHBA) StructureGraph mapped to Uberon. DHBA is implemented within the NIMP Portal for assigning anatomical labels to specimen locations. | |
| **Experimental Factor Ontology** | RRID:SCR_003574 | A systematic description of many experimental variables available in EBI databases. It combines parts of several biological ontologies, such as UBERON anatomy, ChEBI chemical compounds, and Cell Ontology. The scope of EFO is to support the annotation, analysis and visualization of data handled by many groups at the EBI and as the core ontology for Open Targets. | |
| **Harmonized Ontology of Mammalian Brain Anatomy (HOMBA)** | RRID:SCR_027628 | Harmonized cross-species taxonomy of 2,341 brain and spinal cord structures. Derived from the Allen Developing Human Brain Atlas (DHBA) ontology, the HOMBA is hierarchical, allowing users to aggregate structures from fine grain parcellations to broad regions. Terminology is harmonized across human, primate, and rodent structures. | |
| **Ontology of Biomedical Investigations** | RRID:SCR_006266 | An integrated ontology for the description of life-science and clinical investigations. | |
| **UBERON Ontology** | RRID:SCR_010668 | An integrated cross-species anatomy ontology covering animals and bridging multiple species-specific ontologies. | |
| **Metadata** | | | |
| **BICAN Knowledge Base LinkML Schemas** | — | Brain data models in LinkML for the development of the Brain Cell Knowledge Base (BCKB) to ingest and standardize comprehensive cell type information from BICAN's multimodal, multi-species brain cell atlas. | |
| **BICAN Metadata Standards** | 10.5281/zenodo.19377944 | Full suite of BICAN metadata standards hosted in GitHub. | |
| **Data Formats** | | | |
| **AnnData** | RRID:SCR_018209 | A Python package for handling annotated data matrices in memory and on disk. | |

| Resource | RRID / PID | Description | Usage Notes |
|---|---|---|---|
| **Brain Imaging Data Standard (BIDS)** | RRID:SCR_016124 | Provides a consistent way to organize neuroimaging data. | |
| **Loom** | RRID:SCR_028253 | File format designed to efficiently hold large omics datasets. Typically, such data takes the form of a large matrix of numbers, along with metadata for the rows and columns. Developed by the Linnarson Lab. | |
| **Neurodata Without Borders (NWB 2.0)** | RRID:SCR_015242 | A data standard for neurophysiology to share, archive, use, and build analysis tools for neurophysiology data. | |
| **OME-Zarr** | — | A cloud-optimized bioimaging file format. | |
| **SWC** | RRID:SCR_027631 | Text-based (ASCII) files that describe three-dimensional neuronal or glial morphology as a vectorized tree structure of connected nodes. | |
| **Tools** | | | |
| **Annotation Comparison Explorer** | RRID:SCR_026496 | Web application for comparing cell type assignments and other cell-based annotations (e.g., donor demographics, anatomic locations, batch variables, and quality control metrics). Used for connecting brain cell types across studies of health and Alzheimer's Disease. | |
| **Azimuth** | RRID:SCR_021084 | Web application that uses an annotated reference dataset to automate the processing, analysis, and interpretation of a new single-cell RNA-seq or ATAC-seq experiment. | |
| **Basal Ganglia Epigenome Portal** | RRID:SCR_027887 | An integrative resource for exploring epigenetic regulation in the human brain through a web-based platform that unifies single-cell transcriptomic and epigenomic data from the basal ganglia. | |
| **BICAN Experimental Protocols** | — | Collection of protocols developed by BICAN and BICCN available through protocols.io. | |
| **BICAN Human Open Access License (BICAN-BY-NR)** | — | License that explicitly forbids attempts to re-identify individual human research participants from whom the data were obtained. Re-identification includes using these data alone or in combination with other data sets and metadata to identify participants in a de-identified dataset. | *Used for BICAN post-natal human data consented for open release* |
| **BRAIN Armamentarium AAV Collection** | RRID:SCR_027885 | A collection of molecular genetic reagents to gain access to many different brain cell types, available via AddGene. | *Hunker et al. 2025 for striatal focused tools* |
| **Brain Cell Data Viewer** | RRID:SCR_023321 | Web application to view and explore brain cell data with interlinked tabs for Slide-seq atlas gene expression, snRNA-seq atlas, and spatial localization of cell types. | |
| **BRAINCELL-AID** | RRID:SCR_027398 | A comprehensive resource for mouse brain cell cluster annotations, integrating marker gene sets and large language model (LLM) for literature-grounded interpretations across 5,322 brain cell clusters. | |

| Resource | RRID / PID | Description | Usage Notes |
|---|---|---|---|
| **Cell Annotation Platform (CAP)** | RRID:SCR_022797 | A community-driven platform to create, explore, and store cell annotations with associated molecular signatures to interpret cellular identities and encourage convergence upon consensus nomenclature. | *HMBA Consensus Basal Ganglia Atlas available through CAP* |
| **Cellarium Cell Annotation Service (CAS)** | RRID:SCR_025438 | Resource for integrated cell type annotation co-developed by 10X Genomics and Broad Institute Data Sciences Platform. CAS builds a vector search index derived from low-dimensional embeddings of a comprehensive repository of publicly available single-cell transcriptomics data, and uses machine learning algorithms for rapid and accurate annotation of single-cell omics data. | *Williams et al. (2025)* |
| **Cross-enhancer Ranking Pipeline (CERP)** | RRID:SCR_027947 | General peak calling pipeline for multiple species that outputs peak lists per cell type for annotation and filtering. | |
| **Cytosplore Viewer** | RRID:SCR_018330 | An interactive visual analytics system for exploration of single cell and spatial transcriptomics data published in the Allen Cell Types Database and BICAN resources, including the HMBA and 2025 Basal Ganglia release. | *HMBA Human Atlas and basal ganglia data in Cytosplore* |
| **CZ CELLxGENE Discover** | RRID:SCR_024894 | A free, open-source platform from the Chan Zuckerberg Initiative for searching, visualizing, and analyzing large-scale, standardized single-cell transcriptomics data. | |
| **Genetic Tools Atlas** | RRID:SCR_025643 | Searchable web tool for information on enhancer-AAVs and mouse transgenes developed at the Allen Institute, offering a large genetic toolkit for selective gene expression in brain cell types of interest. | |
| **Human Connectome Workbench** | RRID:SCR_008750 | An open source visualization and discovery tool used to map neuroimaging data, especially data generated by the Human Connectome Project. | |
| **MapMyCells** | RRID:SCR_024672 | Maps single cell and spatial transcriptomics data sets to massive, high-quality cell type taxonomies, enabling the creation of brain reference atlases. Supports up to 327 million cell-gene pairs. | |
| **NeMO Analytics** | RRID:SCR_018164 | A web-based, interactive platform designed for visualizing and analyzing complex single-cell transcriptomic and epigenomic data. | |
| **Neuroglancer** | RRID:SCR_015631 | A high-performance, web-based viewer designed for the visualization of large-scale, 3D volumetric data, particularly in the fields of neuroscience and connectomics. | *HMBA Common Coordinate Spaces in Neuroglancer* |
| **Slide-Tags Viewer** | RRID:SCR_028255 | Online viewer for human striatal slide-tags spatial transcriptomics (10X Chromium 5' GEX) dataset comprising ~1.1 million cells from 19 human striatum donors from the Macosko Lab, Broad Institute of Harvard and MIT. | *Kraft et al. (2026)* |
